# Action Progress Provides an Abstract Coordinate for Motor Memory

**DOI:** 10.64898/2026.02.09.704807

**Authors:** Yuto Makino, Kento Suemitsu, Masaya Hirashima

## Abstract

Many everyday actions, such as handwriting, can be performed fluently across a wide range of scales and speeds. This flexibility suggests that motor behavior relies on abstract representations that transcend low-level motor or sensory variables. However, how such abstract representations are formed from individual movement experiences remains unclear. Here, we test the hypothesis that motor memory is organized not only by physical state variables, but also by an abstract coordinate of action progress: the normalized progression of an intended action from initiation to completion. Using a series of motor adaptation experiments that dissociate action progress from kinematic variables and absolute time, we show that learned outputs are retrieved according to their position within an action. This progress-based coordinate spans the unfolding course of a planned action, whether a single reach or an action sequence, and operates continuously rather than as a simple early–late distinction. Together, these findings identify action progress as an abstract organizing coordinate through which the motor system may encode movement experiences as abstract motor memories, offering a framework for understanding the flexible generalization of human motor behavior.

## Introduction

The idea that skilled actions depend on abstract motor representations has a long history in motor control research. Skilled behaviors often preserve a core spatiotemporal structure despite changes in movement scale or speed, a property known as scale–time invariance (1–3). For example, handwriting exhibits consistent letter geometry and characteristic velocity profiles across changes in size and speed (4), and gait shows stable relative timing across walking speeds (5). These observations have motivated classical accounts such as generalized motor programs and schema theory, which propose that skilled actions preserve invariant structural features while allowing execution parameters—such as timing, force, or scale—to vary across contexts (5,6).

More recently, motor abstraction has attracted renewed attention, as this classical idea is now being tested as a functional and neural mechanism for learning and generalization. Behavioral studies have shown that abstract-level motor training can causally support the early stages of learning novel movement patterns (7). In parallel, neurophysiological work has identified action symbols—recombinable representations of discrete units of motor behavior—in the ventral premotor cortex (8). Together, these findings suggest that abstract motor representations are not merely theoretical constructs for explaining well-learned skills, but functional components of the motor system that actively support learning and generalization.

However, a key question remains: how such abstract, generalizable representations are formed from individual movement experiences. Here, we address this representational gap by proposing that motor memory is not merely tied to the features of individual movements, but also organized by action progress—an abstract internal coordinate that spans the unfolding course of an intended action, from initiation to completion.

From a control perspective, the framework of dynamical movement primitives proposes that expressing control commands as a function of a normalized progression variable—from movement onset to offset—enables faithful reproduction and robust generalization of discrete movements across scales (9). Consistent with this view, neural population activity has been shown to evolve along similar trajectories from movement onset to offset, largely independent of movement speed or position (10, 11). Motivated by this convergence, we reasoned that the relative position along such trajectories—namely, action progress—may provide a natural coordinate for organizing abstract motor memory.

To test this representational hypothesis, we turned to motor adaptation, where patterns of adaptation and their generalization provide a window into the structure of the underlying memory (12). A large body of research—particularly studies using force-field adaptation paradigms—has emphasized that motor adaptation is highly specific to the individual movement features experienced during learning (13–18). A widely held view is that adaptation is supported by motor primitives tuned to limb state variables such as position and velocity: learning acquired in a specific state is expressed most strongly when the system revisits nearby states and decays as the state diverges. Although this framework provides a clear computational account of local generalization around trained states (13,19), it remains unclear whether movement state alone is sufficient to explain how motor memories are organized into abstract forms that support broader generalization across actions. In this sense, action progress is not intended as an alternative to state-based organization, but as a higher-order coordinate that may operate alongside physical state variables.

We tested this hypothesis by asking whether motor memories are expressed according to action progress, how this progress coordinate is defined over a planned action, and whether it functions as a continuous dimension. If action progress provides an abstract coordinate for motor memory, learned outputs should be retrieved according to their relative location within an action, even when physical state variables or absolute time differ. Moreover, the coordinate’s relevant range should also be defined not by a single movement element alone, but by the planned action unit. We first tested progress-dependent generalization within a single discrete reach, then asked whether the coordinate can span a double-reach sequence separated by a brief pause. Finally, using a button–reach–button sequence that allowed reach timing to be treated parametrically, we tested whether motor memories are organized within a continuous progress coordinate and whether its boundaries are defined by the planned sequence as a whole.

## Results

### Generalization across reaching distances within a single movement

Experiment 1 examined how motor memory acquired within a single point-to-point reaching movement is generalized to reaches of other distances. Typical motor adaptation studies utilize a curl force field in which force is applied in a consistent direction throughout the single reach. In contrast, the present study used an S-shaped force field with the direction reversed at the midpoint of the reach (14, 20), which requires participants to learn both leftward and rightward compensatory forces within a single reach. This design allowed us to determine where within the movement each learned force component was retrieved and thereby test whether motor memory is expressed along an action-progress coordinate within a single movement.

Participants (n = 16) moved a cursor to a visual target using a manipulandum. They were randomly assigned to one of two groups: Group 1 practiced reaches of 8 cm (Fig. 1a; n = 8), and Group 2 practiced reaches of 16 cm (Fig. 1b; n = 8). During the learning session, participants adapted to the S-shaped perturbation, the magnitude of which gradually increased over trials. The subsequent test session examined how the learned motor memory generalized to an untrained reaching distance. To do so, probe trials were interleaved once every five perturbation trials. For each probe trial, the hand trajectory was constrained to a straight path between the start position and the target using a force channel (clamp channel). The lateral force exerted against the channel wall was used as an index of the retrieved motor memory (21).

**Figure 1.**
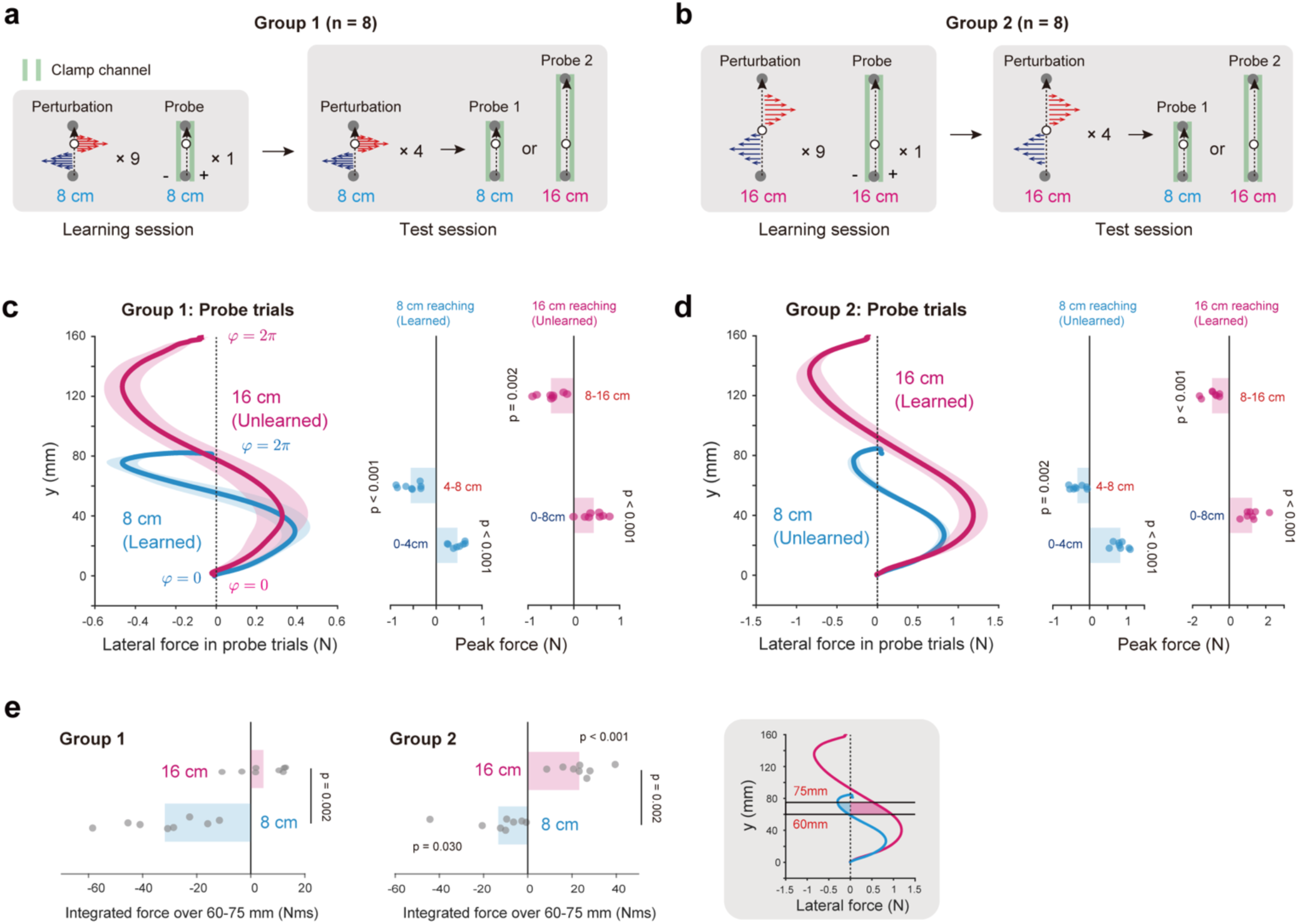
State-dependent representations cannot simply explain scale-invariant generalization in a single reach. **a,b,** Experimental procedures. In Group 1 (a), participants adapted to a position-dependent force field during an 8-cm reaching movement, with the force direction reversed at the midpoint of the movement (4 cm). An 8-cm probe trial was interleaved every ten trials to assess learning. After adaptation, test blocks consisted of four force-field trials followed by either an 8-cm or 16-cm probe trial. Learning-related force output was measured as the force exerted against a clamp channel that constrained the hand to a straight path. In Group 2 (b), participants adapted to an analogous position-dependent force field during a 16-cm reaching movement, with the force direction reversed at the midpoint of the movement (8 cm). Generalization to the untrained distance was assessed as in Group 1. **c,** Forces exerted by Group 1 during probe trials for the two movement distances, plotted as a function of hand position (left). Peak force in the direction opposite to the imposed force field was calculated separately for the first and second halves of the 8-cm and 16-cm reaches (right). Participants expressed the appropriate force direction in both the trained 8-cm reach (0–4 cm, t(7) = 8.253, p < 0.001; 4–8 cm, t(7) = 7.146, p < 0.001) and the untrained 16-cm reach (0–8 cm, t(7) = 4.618, p = 0.002; 8–16 cm, t(7) = −5.515, p < 0.001). φ indicates reach progression. **d,** Forces and peak forces for Group 2. Participants expressed the appropriate force direction in both the trained 16-cm reach (0–8 cm, t(7) = 7.525, p < 0.001; 8–16 cm, t(7) = −6.294, p < 0.001) and the untrained 8-cm reach (0–4 cm, t(7) = 12.299, p < 0.001; 4–8 cm, t(7) = −4.791, p = 0.002). **e,** Force impulse was computed by integrating force over the 60–75 mm position range to quantify the force direction expressed at that location. Force magnitude differed significantly between movement distances in both groups.

Figures 1c and 1d present the retrieved force profiles measured during the test session for both learned and unlearned reaching distances. For learned distances, participants expressed clear compensatory force components in opposite directions in the first and second halves of the movement, as required to counteract the S-shaped perturbation. The peak forces in the first and second halves of the movement were significantly directed opposite to the corresponding force component of the perturbation (t-tests, p < 0.001; learned panels in Fig. 1c–d). Importantly, a similar force pattern was observed when participants performed the unlearned distance.

Even though the reaching distance changed, the retrieved force again reversed between the first and second halves of the movement, and the peak forces in the first and second halves of the movement were significantly directed opposite to the perturbation (t-tests, all p < 0.002; unlearned panel in Fig. 1c–d). Thus, the learned force pattern generalized across reaching distances while preserving its relative organization within the movement.

This pattern cannot be explained by a purely velocity-dependent account. The first and second halves of the reach involved nearly identical hand velocities, yet participants retrieved opposing force components in these two halves. If motor memory were indexed solely by velocity, the memories acquired in the two halves should have interfered with each other or been retrieved similarly. Instead, participants generated appropriately opposite forces in the two halves of the movement, indicating that velocity alone cannot account for observed memory expression.

A position-dependent account was also insufficient. If the two opposing force components were tied to absolute limb position, then the same compensatory force should be retrieved whenever the hand passes through the same spatial location, regardless of the reach distance. However, this was not observed. For example, in Group 2, the first 8 cm of the trained 16-cm reach exhibited a predominantly rightward compensatory force. However, when the same participants performed the untrained 8-cm reach, the force reversed within that same spatial region, with rightward force in the first half and a leftward force in the second half (see the force profile for the unlearned distance in Fig. 1d). Thus, at overlapping limb positions, the retrieved force differed depending on whether that position occurred in the first or the second half of the movement. To quantify this discrepancy, we compared the lateral force between the learned and unlearned movements within a shared spatial window of 60–75 mm (Fig. 1e inset), corresponding to the second half of the 8-cm reach and the first half of the 16-cm reach. The retrieved force differed significantly between reaching distances in both groups (Student’s t-tests: Group 1, t(7) = 5.062, p = 0.002; Group 2, t(7) = 4.866, p = 0.002; Fig. 1e), further arguing against position-dependent retrieval alone.

Taken together, these results show that neither position-dependent nor velocity-dependent encoding alone is sufficient to explain the generalization observed in Experiment 1. Additional analyses further showed that accounts based on other individual physical variables were also insufficient. The force profiles could not be explained by memory tied to absolute elapsed time: when trials were divided into fast and slow subsets based on the time taken to reach the movement midpoint, the force profiles did not follow a fixed temporal pattern but instead were better aligned after temporal scaling (Supplementary Fig. 1). This pattern is consistent with force retrieval following the relative progress of the movement, rather than absolute elapsed time. The observed force profiles were also inconsistent with an account based solely on acceleration magnitude (Supplementary Fig. 2). Thus, the findings are more parsimoniously explained by a relative coordinate, such as progression through the reach, rather than by any single absolute-valued kinematic or temporal variable.

### Progression-based organization in double-reach sequences

Experiment 2 asked whether the progress coordinate can span an entire sequence when two reaches are planned as a unified action, using a double-reach task in which participants briefly stopped at an intermediate point (Fig. 2a). If each reach is planned independently, motor memory should be organized within separate progression cycles for each component reach. However, if two reaches are planned as a unified action, motor memory should instead be organized over a single progression cycle spanning the entire sequence.

**Figure 2.**
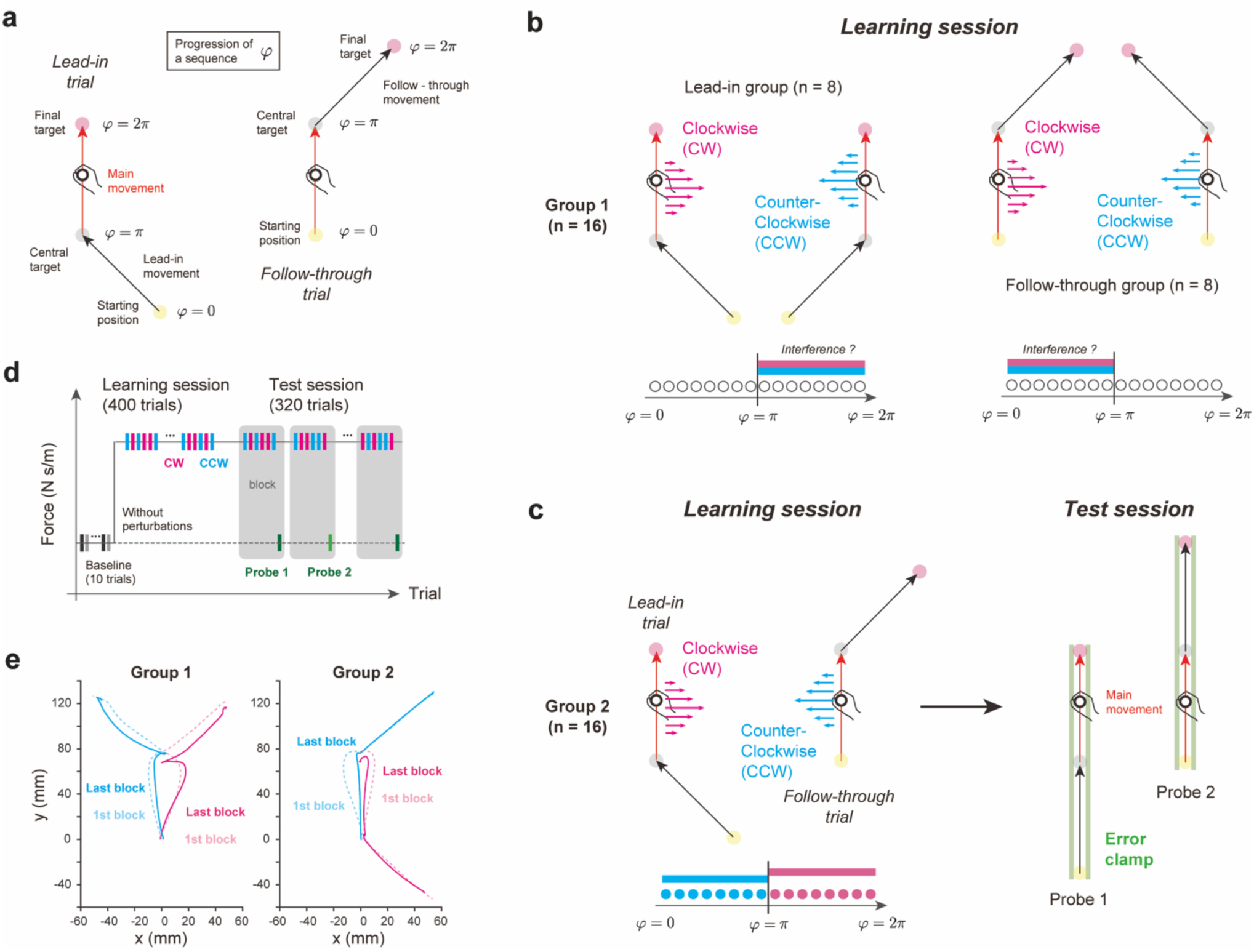
Progression-dependent learning during the double-reaching movement. **a,** Progression was defined over the double-reach sequence such that the start, intermediate stop, and endpoint corresponded to 0, π and 2π, respectively. **b,** Experimental paradigm motivated by previous studies (Howard et al., 2012, 2015). In these studies, participants slowly learned different force fields during an identical main reach depending on the direction of the preceding lead-in or subsequent follow-through movement (top). Under a progression-dependent memory framework, opposing force fields should interfere when they are assigned to the same progression range, leading to slower learning (bottom). Group 1 was trained in this conflicting force-field condition. **c,** Force-field condition learned by Group 2. The direction of the force field varied according to trial type (lead-in or follow-through trial; top left), such that the perturbed reach occupied different progression ranges within the double-reach sequence (bottom left). After training, two types of probe trials were interleaved to assess memory retrieval (right). In these probe trials, the tested reach corresponded to either the first or second movement of the sequence. **d,** Experimental timeline. During test sessions, probe trials were interleaved every seven trials. **e,** Hand trajectories from representative participants in Groups 1 (left) and 2 (right). In the first block of ten force-field trials, trajectories were markedly perturbed by the force field (dotted lines). In the final blocks, participants adapted and produced nearly straight movements toward the target (solid lines).

Previous studies have shown that participants can learn opposing curl force fields when each force field is associated with the direction of a lead-in movement performed immediately before the main reach (22) or a follow-through movement performed immediately after it (23) (Fig. 2b). These contextual effects are typically interpreted as showing that different preceding or subsequent movements cue the retrieval of distinct motor memories (22, 23). Here, we reinterpret those findings from the perspective of action progress. If the entire double-reach sequence is represented as a single unified action, then opposite force fields assigned to the same portion of the sequence should interfere, whereas force fields assigned to different portions of the sequence should be mapped to different locations along a sequence-level progress coordinate and should therefore interfere less.

To test this idea, we assigned 32 participants to two groups (Fig. 2b-c). During the learning session (Fig. 2d), Group 1 participants (Fig. 2b; n = 16) performed either lead-in trials or follow-through trials in which opposite force fields were associated with two different directions of the same contextual movement type, meaning that the opposing perturbations were experienced within the same progression range of the sequence (high-interference condition). Group 2 participants (Fig. 2c; n = 16) performed both lead-in and follow-through trials, and opposite force fields were associated with different trial types, meaning that the perturbations were experienced in different progression ranges (reduced-interference condition). Thus, the critical distinction between the groups was whether the two opposing force fields occupied overlapping or separated portions of a putative sequence-level progress coordinate.

Group 1 participants exhibited large deviations from a straight path throughout learning, whereas Group 2 participants showed gradual compensation for the perturbations and achieved straighter movements (Fig. 2e). Figure 3a presents the learning curves for the opposing force fields in each group. To quantify learning, we measured the magnitude of lateral deviation at the peak velocity during the perturbed main movement. As predicted by the action-progress framework, Group 2 participants—who experienced the two force fields in different progression ranges—showed facilitated adaptation compared with Group 1 participants, for whom both force fields were associated with the same progression range.

**Figure 3.**
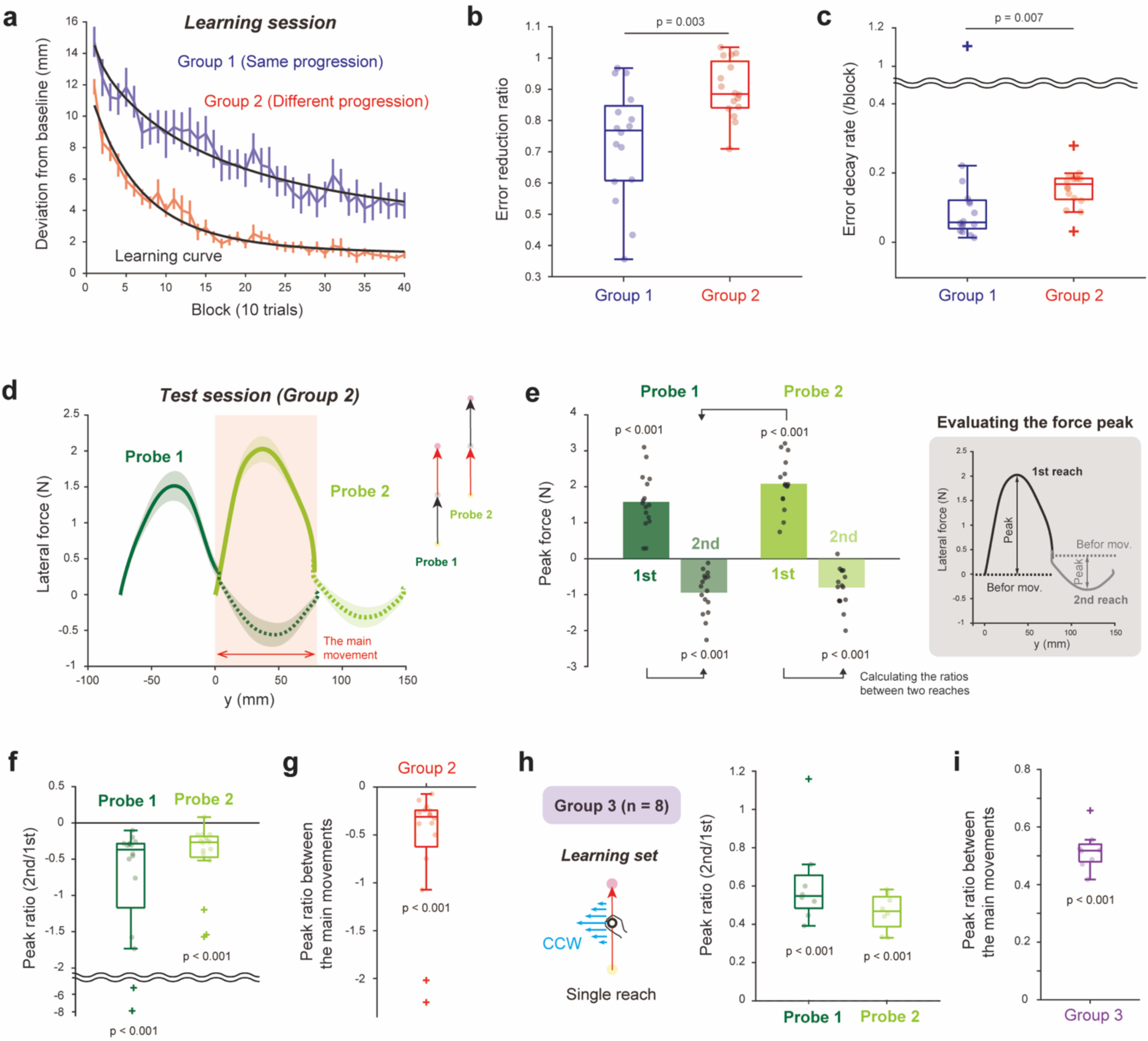
Motor memory is associated with the progression of the double reach. **a**, Learning traces for Group 1 (same progression, blue) and Group 2 (different progression, red) participants. The degree of learning was quantified as the lateral deviation from the straight path at the time of peak velocity. Each trace was fitted with an exponential curve (learning curve; black line). **b,** Comparison of the error reduction ratio between Groups 1 and 2. Group 2 showed a significantly greater decrease than Group 1. **c,** Error decay rate from the exponential fit. **d,** Force exerted to the clamp channel in probe trials. e, Direction of the retrieved motor memory in the probe reaches, calculated as the difference between the peak force during movement and the holding force exerted before movement (inset). Participants exerted significantly positive forces in the first reach (one-sample t-test; Probe 1: t(15) = 8.016, p < 0.001; Probe 2: t(15) = 11.768, p < 0.001) and significantly negative forces in the second reach (one-sample t-test; Probe 1: t(15) = −6.307, p < 0.001; Probe 2: t(15) = −5.884, p < 0.001). **f,** Peak ratios, defined as the peak force during the second reach divided by that during the first reach, indicated a reversal of force direction within the same movement sequence. **g,** Peak ratios, calculated as the peak force of the second reach in Probe 1 divided by that of the first reach in Probe 2 (panel e), demonstrate retrieval of opposite motor memories despite identical movement states. **h,** A control experiment (Group 3) was conducted to test whether the results (d–g) reflected progression-dependent encoding of distinct motor memories. Participants learned a force field during a single reach and were then tested during a double forward reach, as in Group 2. No reversal of the retrieved force was observed between the first and second reaches. **i,** Consistent with the findings for Group 2, Group 3 participants exerted lateral forces in the same direction during the second reach in Probe 1 and the first reach in Probe 2.

We evaluated group differences in error reduction using two complementary metrics. First, the overall reduction in error, quantified by the error reduction ratio (1 −*E_last_*/*E_first_*), where *E_first_* and *E_last_* denote the mean lateral deviation in the first and last ten trials, respectively, was significantly greater in Group 2 than in Group 1 (Mann–Whitney U test, z = −2.959, p = 0.003; Fig. 3b). Second, the exponential error decay rate obtained by fitting *y* = *Ae*^−*Bx*^ + *C* to the trial-by-trial time course of lateral deviation was also significantly larger in Group 2 than in Group 1 (Mann–Whitney U test, z = −2.695, p = 0.007; Fig. 3c). Even when restricting the comparison to the lead-in subgroup of Group 1—which is suggested to exhibit relatively rapid learning (22, 23)—Group 2 still demonstrated significantly facilitated adaptation (Supplementary Fig. 3). These results indicate that separating opposing force fields across different portions of the sequence reduces interference.

Although Group 2 showed faster learning than Group 1, this group difference alone does not establish progression-dependent organization. One possible interpretation is that assigning the two force fields to different progression ranges reduced interference between the corresponding memories. However, Group 2 also had both the preceding and following movements available as potential contextual cues, whereas each subgroup of Group 1 had only one contextual movement type. Thus, improved adaptation could in principle reflect the availability of multiple cues rather than the organization of memories along action progress. To dissociate these possibilities, we examined memory retrieval in two error-clamp probe conditions (Probe 1 and Probe 2; Fig. 2c) that contained a shared kinematically identical movement (Fig. 2c, red segment) occurring either in the first or second half of the sequence composed of two forward reaches. If the two force fields had been learned at distinct locations along a sequence-level progress coordinate, the force output should reverse between the first and second reaches. Critically, these probes allowed us to test whether motor memories organized by action progression could be retrieved even when state information provided by the immediately preceding and following movement directions was altered.

Figure 3d shows the lateral force profiles measured in the channel trials. In both probes, the force direction was reversed between the first and second reaches. We quantified this reversal by measuring the peak lateral force for each reach relative to its value at reach onset (inset of Fig. 3e). The peak values are summarized in Fig. 3e. Next, we computed the signed ratio of the second-reach peak to the first-reach peak (Fig. 3f), such that negative values indicate opposite-signed forces across the two reaches. This ratio was significantly negative in both Probe 1 (median [interquartile range, IQR] = −0.369 [−1.169, −0.286]; Wilcoxon signed-rank test, p < 0.001) and Probe 2 (median [IQR] = −0.268 [−0.473, −0.186]; Wilcoxon signed-rank test, p < 0.001), confirming that the retrieved force was reversed between the first and second reaches in both probe conditions.

A particularly critical comparison is between the second reach of Probe 1 and the first reach of Probe 2. These two movements are kinematically identical (Supplemental Fig. 4) and occur at the same limb position where the force field had been experienced during learning, differing only in whether the movement occurred in the early or late portion of the double-reach sequence. If motor memory were determined solely by state-dependent variables (e.g., position or velocity), the retrieved forces should be identical. Similarly, if the learning advantage in Group 2 merely reflected the availability of multiple contextual cues, this account would not specifically predict opposite retrieved forces between these kinematically matched reaches. Instead, the measured forces had opposite signs (Fig. 3g), indicating that distinct motor memories are learned and retrieved according to where the movement occurred within the sequence-level progress coordinate.

Finally, we carried out a control experiment to confirm that the force reversal was not simply an artifact of the two-reach error-clamp probe. If the reversal truly reflects progression-dependent learning of opposing force fields, it should be absent when participants learn a single force field. A separate control group (Control Group 3; Supplementary Fig. 5) learned a single force field during a single reach (Fig. 3h, inset) and then performed the same two-reach error-clamp probe trials. In this group, no action-progress reversal was observed. The signed ratio of the second-reach peak force to the first-reach peak force was significantly positive in Probe 1 (median [IQR] = 0.548 [0.483, 0.656]; Wilcoxon signed-rank test, p < 0.001) and Probe 2 (median [IQR] = 0.469 [0.388, 0.544]; Wilcoxon signed-rank test, p = 0.008; Fig. 3h). Likewise, the comparison between the second reach of Probe 1 and the first reach of Probe 2 was significantly positive (median [IQR] = 0.498 [0.421, 0.543]; Wilcoxon signed-rank test, p = 0.008; Fig. 3i), indicating that opposite outputs were not retrieved when only a single memory had been learned.

Overall, these findings suggest that, when two reaches are planned as a unified action, motor memory can be organized according to where a movement occurs within the sequence.

### A parametric approach to the sequence-level organization of motor memory

Experiment 2 showed that motor memory differs between the early and late portions of a double-reach sequence. However, this result leaves open whether memory is organized along a continuous progress coordinate, or whether it simply differs between two discrete contexts: the first and second reaches. Experiment 3 addressed this question using a button–reach–button task (Fig. 4a) in which participants performed a left-hand button press (B_1_), a right-hand reach (R), and a second left-hand button press (B_2_). By controlling the timing of the three-action sequence with a progress bar (Fig. 4b), this task allowed us to vary parametrically the progression value at which the reach occurred within the overall sequence.

**Figure 4.**
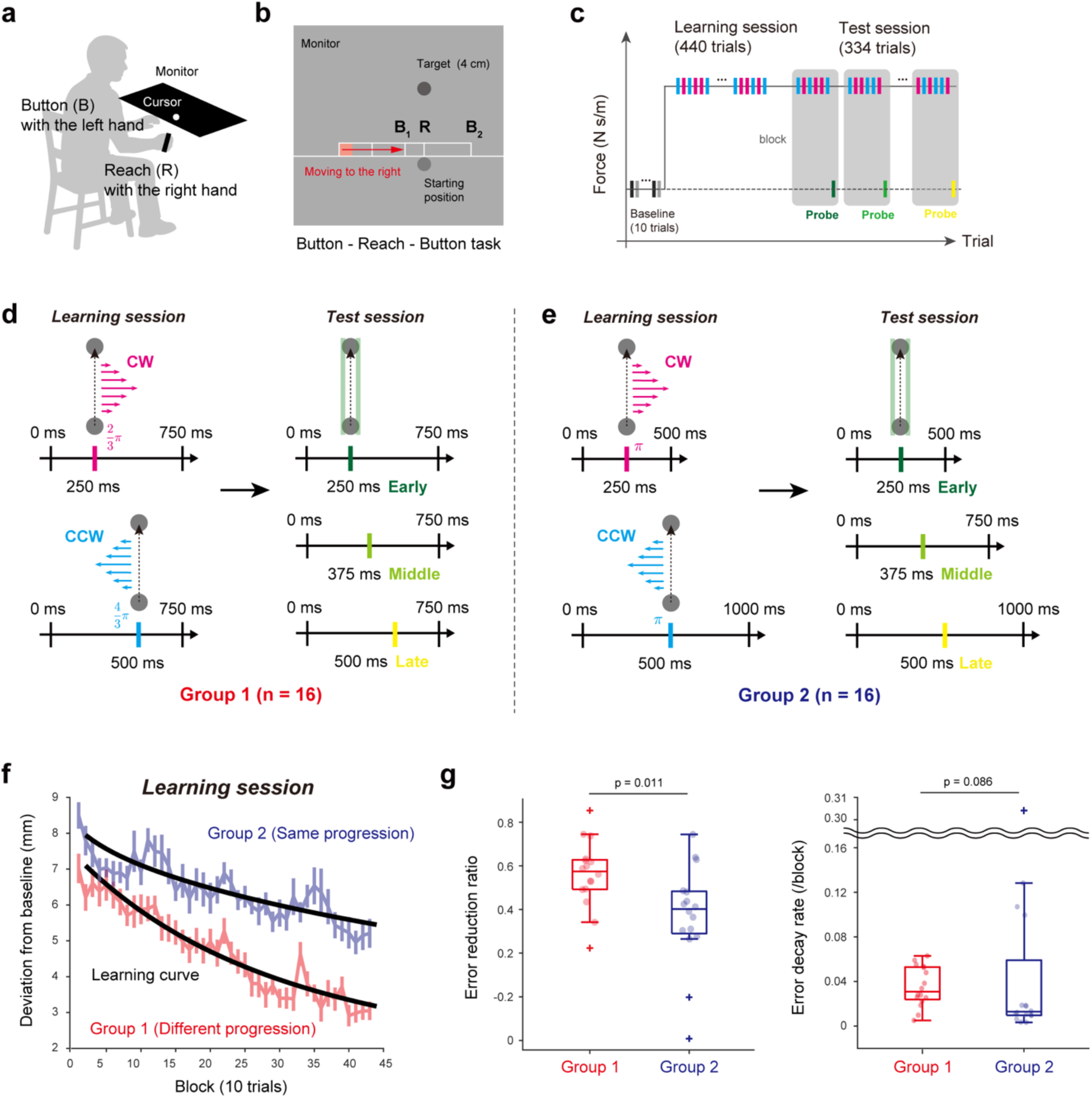
Experimental paradigm for a parametric approach to sequence-level organization of motor memory. **a,** Schematic of Experiment 3. Participants performed reaching movements with their right hand while pressing a button with their left hand before and after each reach. **b,** A visual bar moved from left to right at a constant speed to indicate the timing of the B_1_–R–B_2_ sequence. Participants pressed the button (B_1_) at the third and fifth ticks and performed the reach (R) at the fourth tick. The bar disappeared at the first button press (B_1_), encouraging participants to plan the entire B_1_–R–B_2_ sequence in advance. **c,** Structure of the experimental trial, which consisted of a learning session and a test session as in Experiment 2. **d,** For Group 1, the interval between the two button presses (B_1_–B_2_) was 750 ms. The timing of the reach (R) was varied within this interval. The direction of the force field in the learning session was switched according to the reach timing (progression). After learning, probe trials were conducted under the clamp channel where participants performed reaches at 2π/3, π, and 4π/3 progression of the sequence. **e,** For Group 2, progression of the reach was fixed, and the direction of the force field in the learning session was switched according to the interval between two button presses. In the probe trials, the interval was set to 500, 750, or 1000 ms. **f,** Learning traces for Groups 1 (different progression, red) and 2 (same progression, blue). g, Comparison of the error reduction ratios (left) and error decay rates (right) between Groups 1 and 2.

We first asked whether learning opposing force fields is facilitated when the two force fields are separated across different action-progress values rather than assigned to the same action-progress value. Participants (n = 32) were randomly assigned to two groups. In both groups, the direction of the curl field was associated with one of two B_1_–R intervals (250 or 500 ms), but the mapping from these intervals to progression within the full button–reach–button sequence differed between groups. In Group 1 (Fig. 4d), the total button-to-button duration was fixed at 750 ms such that the reach occurred at different progression values depending on the B_1_–R interval: 2π/3 for the 250-ms condition and 4π/3 for the 500-ms condition. In this group, progression provided an additional cue for force-field direction alongside the absolute B_1_–R interval. In contrast, in Group 2 (Fig. 4e), the total duration was scaled with the B_1_–R interval (500 ms vs. 1000 ms) so that the reach always occurred at the same progression value (π) despite different absolute intervals. Thus, for Group 2, the force-field direction could be predicted only from the absolute timing variables (B_1_–R, R–B_2_, and total duration), not from progression. The CW and CCW force fields were presented in pseudorandom order for both groups.

Learning was facilitated in Group 1 relative to Group 2 (Fig. 4f). The error decay rate did not differ significantly between groups (Mann–Whitney U test, z = 1.715, p = 0.086; left panel in Fig. 4g), whereas the error reduction ratio was significantly larger in Group 1 than in Group 2 (Mann–Whitney U test, z = 2.544, p = 0.011; right panel in Fig. 4g). These results demonstrate that separating opposing force fields across different progression values facilitates learning during training.

We next asked whether the learned memories were expressed as a graded function of the progress coordinate. Probe trials sampled not only the two trained conditions but also an intermediate condition, allowing us to examine how memory expression varied across the coordinate. Figure 5a illustrates the forces exerted in the probe trials. The compensatory force size was quantified as the impulse of the lateral force from movement onset to peak velocity, normalized to the impulse of the ideal force (computed as b⋅v) over the same interval and expressed as a percentage (Fig. 5b). In Group 1, the compensatory force size was systematically modulated by progression (one-way repeated measures analysis of variance (ANOVA), F(2,80) = 184.8, p < 0.001): the two trained conditions elicited opposite compensatory forces, whereas the intermediate condition produced an intermediate level of forces (post hoc comparisons with Holm correction; Early vs. Middle: t(15) = −10.62, p < 0.001; Middle vs. Late: t(15) = −10.94, p < 0.001). This graded pattern indicates that motor memory was not merely assigned to two discrete contexts but was organized along a continuous sequence-level progress coordinate.

**Figure 5.**
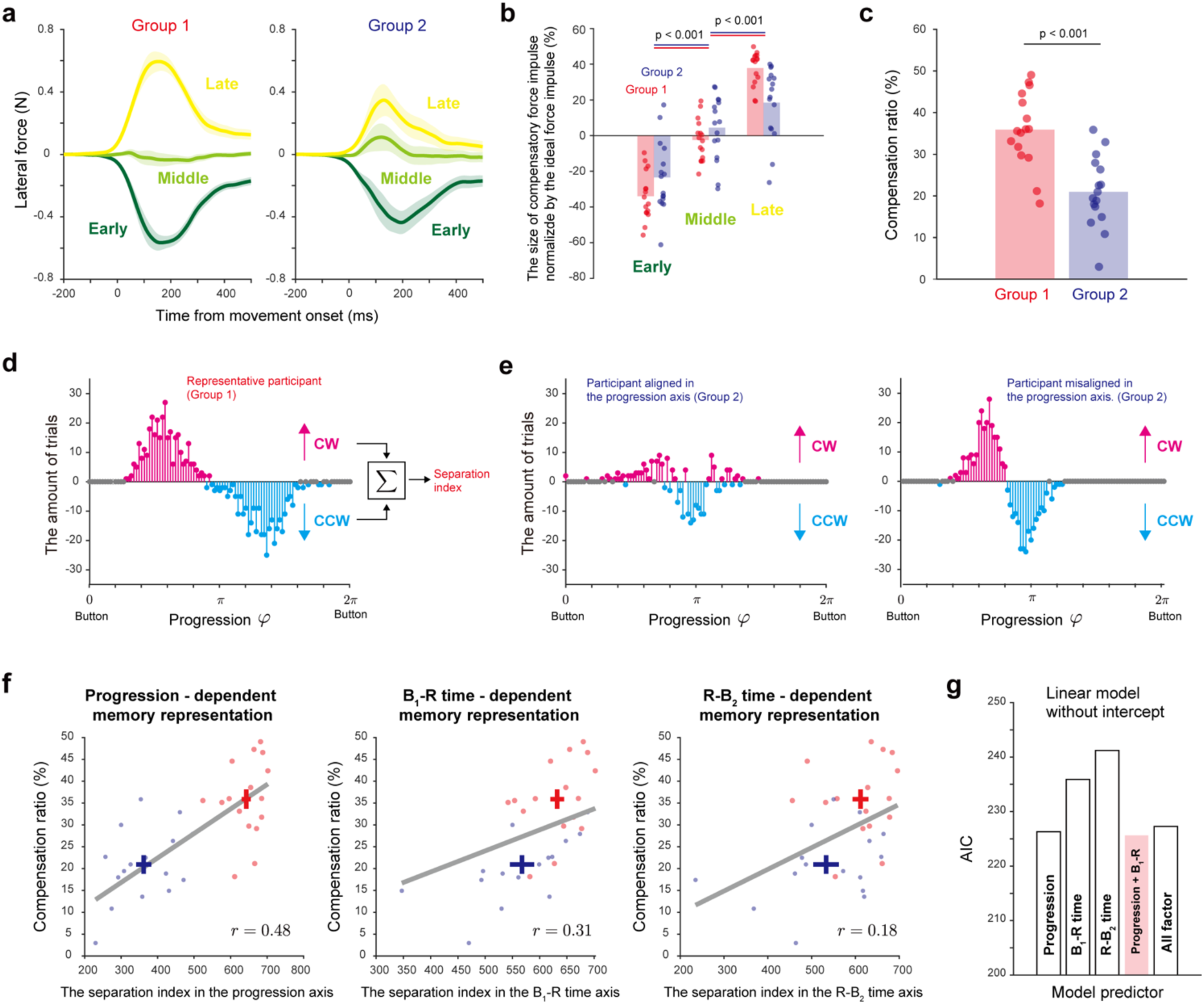
Progression–dependent memory representation for motor learning of a sequence of movements. **a,** Exerted forces in the probe trials for Groups 1 and 2. Participants produced negative forces in the early progression (Group 1) or interval (Group 2), which corresponded to the clockwise (CW) perturbation; positive forces in the late progression (Group 1) or interval (Group 2), which corresponded to the counterclockwise (CCW) perturbation; and forces close to zero in the middle progression value or interval. The overall force magnitude was greater in Group 1 than in Group 2. **b,** Compensatory forces for each sequence pattern were calculated as the impulse of the exerted lateral force from movement onset to peak velocity, normalized by the impulse of the ideal compensatory force (v × b) and expressed as percentages. **c,** Comparison of the total compensation ratios for the two learned sequence patterns between Groups 1 and 2. **d–e**, Quantification of separation along the progression dimension. Progression was divided into 100 bins, summing the experienced force field (–1 for CCW, +1 for CW). Group 1 showed clear separation of memories along the progression axis (d). Some Group 2 participants showed overlapping representations near the middle progression (e, left), whereas others showed progression-separated patterns contrary to the instructions (e, right). Data from a representative participant is shown. **f,** Memory separation along the progression dimension as the best predictor of group differences in learning performance (left). The degree of separation of the two memories along the progression axis explained the group difference in the compensation ratio better than other time-dependent dimensions (middle: B_1_–R absolute time, regression coefficient = 0.048, r^2^= 0.309; right: R– B_2_ absolute time, regression coefficient = 0.050, r^2^= 0.183). **g,** Model comparison using Akaike information criterion (AIC). When comparing models that included two or three temporal factors, the combination of progression and absolute timing between button press and reach (B_1_–R) best explained memory expression.

Group 2 exhibited a qualitatively similar pattern across the trained and intermediate absolute-time conditions (one-way repeated measures ANOVA, F(1.5, 21.75) = 77.19, p < 0.001 with Greenhouse-Geisser correction) with significant differences between adjacent conditions (post hoc comparisons with Holm correction; Early vs. Middle: t(15) = −8.00, p < 0.001; Middle vs. Late: t(15) = −6.10, p < 0.001). However, overall memory expression was stronger in Group 1: the compensation ratio, computed by averaging sign-aligned compensatory forces in the trained conditions, was significantly higher in Group 1 than in Group 2 (Student’s t-test, t(30) = 4.872, p < 0.001; Fig. 5c). Thus, action progress provided a more effective organizing coordinate than absolute timing alone.

Because Group 2 still expressed opposing memories, we next examined whether this reflected true organization by absolute timing or unintended separation along action progress. Even when the task was designed to place the reach at the same progress value in Group 2, participants’ timing variability could shift the realized reach position along the progress coordinate. To examine this possibility, a progression-separation index was computed by binning sequence progression into 100 bins, accumulating the force-field exposures (+1 for CW, −1 for CCW), and summing the absolute values across bins (see Methods, Eq. 3). Higher values indicate greater separation of the two force fields along progression.

Figure 5d shows an example Group 1 participant with a high separation index, reflecting clear segregation of the two fields along progression. In contrast, some Group 2 participants expressed both force fields at nearly the same progression value (Fig. 5e, left), whereas others introduced small but systematic progression shifts that yielded partial separation (Fig. 5e, right). Across participants in both groups, the progression-separation index strongly predicted the degree of memory expression. A linear regression constrained through the origin indicated a positive relationship between the separation index and the compensation ratio (β = 0.056, r^2^ = 0.487; Fig. 5f, left panel).

We then compared progression with absolute temporal intervals as candidate organizing dimensions, as suggested by previous studies (17). Analogous to the progression-based analysis, separation indices along these temporal dimensions were computed by binning each interval into 100 bins per 1 s. The separation indices based on B_1_–R and R–B_2_ intervals explained the compensation ratio less well than the progression-based index (Akaike information criterion (AIC) (progression) = 226.3; AIC(B_1_–R) = 235.9; AIC(R–B_2_) = 241.2; Fig. 5f– g). Model comparison across all combinations of progression, B_1_–R, and R–B_2_ showed that the best-fitting model included progression and the absolute B_1_–R interval (AIC = 225.6), outperforming both the progression-only model (AIC = 226.3) and the model including all temporal variables (AIC = 227.3). These analyses indicate that sequence-level progress was the dominant coordinate for organizing motor memory, although absolute timing from the preceding action (B_1_–R) also contributed.

Finally, we asked how this coordinate is constructed over the planned sequence. Specifically, we compared the button–reach–button task with a simpler button–reach task, in which participants performed only B_1_ and R and opposing force fields were mapped exclusively to the B_1_–R interval (Fig. 6a,b). In this simplified task, the B_1_–R interval was the only dimension for separating memories. Participants showed significant adaptation based on this interval: the error reduction ratio was significantly greater than zero (median [IQR] = 0.389 [0.326, 0.511]; Wilcoxon signed-rank test, z = 3.52, p < 0.001; Fig. 6c) and probe forces differed across B_1_–R intervals during the test session (one-way repeated measures ANOVA followed by post hoc comparisons with Holm correction; Early vs. Middle: t(15) = −6.872, *p* < 0.001; Middle vs. Late: t(45) = −7.525, p < 0.001; Fig. 6d). Thus, absolute timing can serve as an organizing signal.

**Figure 6.**
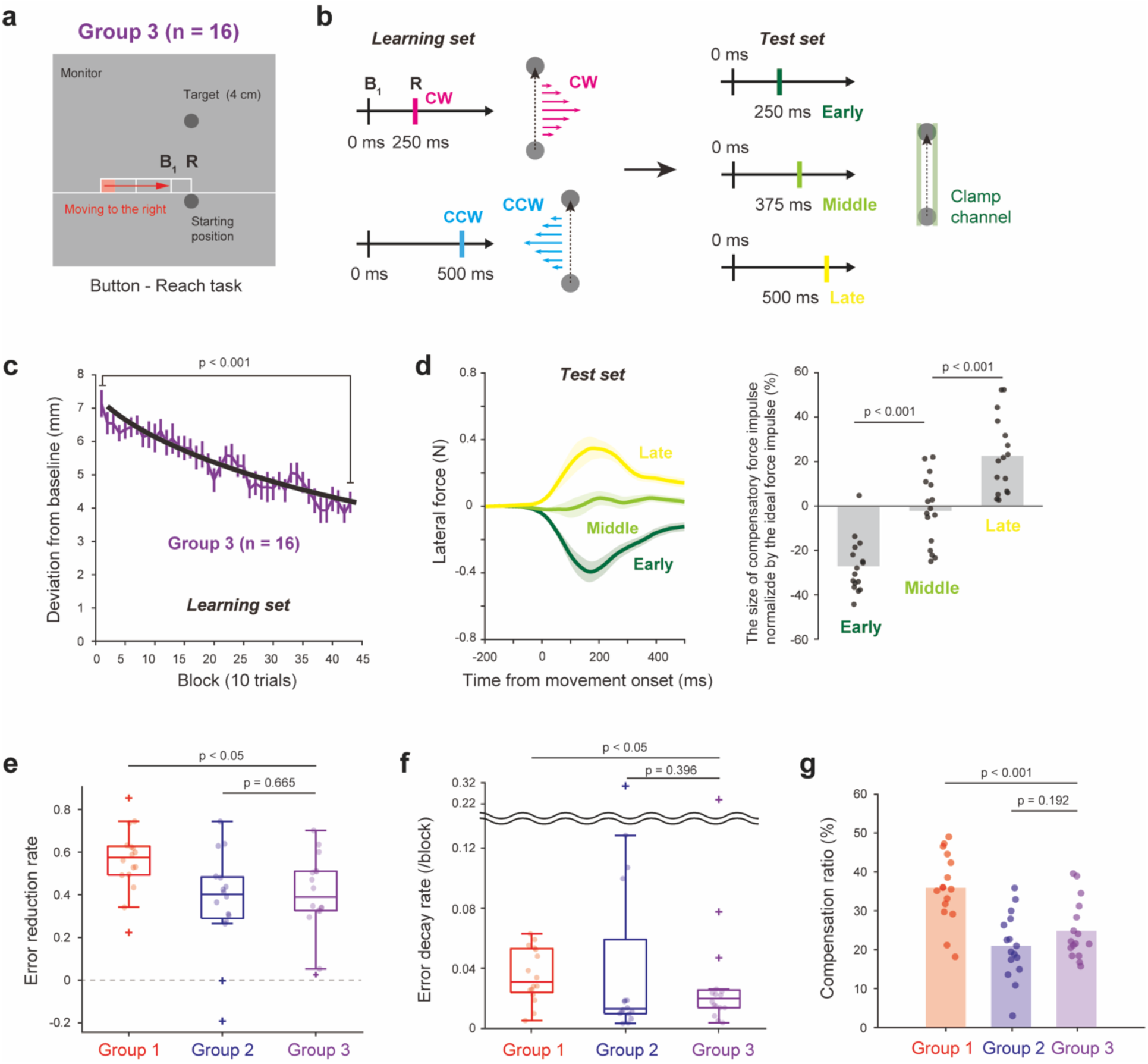
Motor memories are associated with the absolute time from the prior movement, but its effect is weaker than that of progression. **a,** Experimental setup of Group 3. Participants executed a button press followed by a reach (B_1_–R). **b,** Experimental protocol. In the learning set, the direction of the force field was associated with the interval between the two actions. The learning response was evaluated in the probe trials with the interval set to 250, 375, or 500 ms. **c,** Learning traces for Group 3 show that the lateral deviation gradually decreased as the learning set progressed. **d,** Exerted forces in the probe trials (left). Calculation of the size of compensatory forces for each sequence pattern (right). One-way repeated measures analysis of variance (ANOVA) revealed that participants produced appropriate forces according to the time interval (F(2,30) = 96.818, p < 0.001). **e,** Comparison of the error reduction ratio among the three groups. The Kruskal–Wallis test revealed a significant difference in the error reduction ratio among the three groups (H(2) = 8.577, *p* = 0.016). **f,** Comparison of the error decay rate among the three groups. Only a non-significant trend toward a group effect was present. **g,** Comparison of the compensation ratios among the three groups. One-way repeated measures ANOVA revealed a significant difference in the learning ratio among the three groups (F(2,45) = 13.900, *p* < 0.001).

However, its effect was limited. Error-reduction in the control group (Group 3) was lower than that in Group 1 and comparable to that in Group 2, as shown by planned comparisons for the error reduction ratio (Kruskal–Wallis test followed by the Wilcoxon rank-sum test with Holm correction; Group 1 vs Group 3: z = 2.431, p = 0.030; Group 2 vs Group 3: z = −0.433, p = 0.665; Fig. 6e). A similar pattern was observed for the error decay rate: Group 3 was lower than Group 1 but comparable to Group 2 (Group 1 vs Group 3: z = 2.016, p = 0.044; Group 2 vs Group 3: z = −0.848, p = 0.396; Fig. 6f), although the main effect did not reach statistical significance in the error decay rate (H(2) = 5.103, p = 0.082). The force-based compensation ratio showed the same pattern, with Group 3 significantly lower than Group 1 but not different from Group 2 (one-way ANOVA followed by post hoc comparisons; Group 1 vs. Group 3: t(45) = 3.76, *p* < 0.001; Group 2 vs. Group 3: t(45) = −1.33, p = 0.192; Fig. 6g).

Overall, Experiment 3 shows that motor memory can be organized within a continuous progress coordinate. Moreover, the comparison between the button–reach–button and button–reach tasks indicates that this coordinate is defined by the structure of the sequence as a whole, rather than by the absolute timing between adjacent action elements alone. Absolute B_1_–R timing can contribute to memory organization, but it provides a weaker organizing signal than sequence-level action progress.

## Discussion

A hallmark of the motor system is its ability to flexibly reuse motor memories acquired from previous movement experiences to meet new behavioral demands. This flexibility suggests that motor memories are not stored merely as responses tied to the specific kinematics or timing of individual movements but can also be organized within a more abstract representational structure. However, it remains unclear what abstract coordinate system supports this form of organization. Here, using motor adaptation experiments, we provide converging behavioral evidence that action progress can serve as an abstract, continuous coordinate for organizing motor memory within the structure of a planned action.

### Action progress as an abstract coordinate for motor memory

Motor memories are often described as being supported by primitives tuned to limb state variables, such as position and velocity, producing state-dependent generalization (13–18). This framework has been powerful for explaining force-field adaptation, including local generalization and interference when two perturbations overlap in state space. Here, our aim was not to challenge the importance of physical-state coordinates, but to test whether they are sufficient to explain generalization across changes in movement scale.

Experiment 1 provided this test by asking whether a learned force pattern would transfer across reaching distances. Participants learned to compensate for an S-shaped force field and expressed a similar S-shaped force pattern even when reaching over distances different from those experienced during learning (Figs. 1c-d). The acquisition of an S-shaped pattern argues against a purely velocity-based account, because participants expressed oppositely directed compensatory forces within a single reach despite comparable velocity magnitudes across the two movement halves (14). The transfer across reaching distances further showed that force expression at overlapping spatial positions depended on where those positions occurred within the progression of the action (Fig. 1e), indicating that position alone was also insufficient. Additional analyses showed that neither acceleration magnitude nor absolute elapsed time alone provided a sufficient explanation of the observed force profiles (Supplementary Figs. 1–2).

We cannot rule out more complex alternatives in which multiple physical variables, or model-free updates of motor policy (24), account for aspects of the data. However, such accounts would require the motor system to flexibly combine and reweight these variables across changes in movement scale. It is unclear how this strategy would naturally give rise to the generalization observed here. These considerations motivate an action-progress account, in which the learned force pattern is organized not only by physical movement variables, but also according to progression within the intended action, allowing it to be retrieved according to the same progression in a new movement situation.

A related alternative is a gain- or scaling-based account, in which the learned relationship between movement state and motor output is rescaled according to changes in movement distance or speed (25). In the present case, such an account could describe the transfer across reaching distances as a rescaling of the learned force pattern from the trained movement distance to the test movement distance. However, this account leaves open how the motor system selects which previously learned state-dependent memory should guide the current movement and, within that memory, how the relevant part should be aligned and rescaled to the current movement. A progress-based account avoids this selection-and-alignment problem by allowing learned outputs to be read out according to the movement’s relative progress within the intended action, rather than by requiring reference to a particular previously learned state-specific memory. Thus, although state-dependent and gain- or scaling-based mechanisms may contribute to the observed transfer, the action-progress account provides a parsimonious way to describe the component of generalization that depends on where the movement lies within the intended action.

### Sequence-level progression

If action progress provides an organizing coordinate for motor memory, it should not be limited to the internal progression of a single reach, but may also apply to multiple movement elements organized as a single planned action. Experiment 2 tested this possibility using a double-reach task (22–23) in which both the first and second targets were presented before movement onset, allowing participants to know the final goal of the sequence in advance (Fig. 2a–c).

Learning was facilitated when opposing force fields were assigned to different portions of the sequence, but impaired when both fields were experienced within the same portion (Fig. 3b–c). Although this learning advantage could partly reflect the availability of both preceding and following movements as contextual cues in Group 2, the subsequent probe trials provided a more direct test: kinematically identical reaches elicited opposite force outputs depending on whether they occurred early or late within the sequence (Fig. 3d–g). This result provides stronger evidence for progression-dependent retrieval at the sequence level.

This interpretation does not imply that any series of movements is automatically organized into a single progression cycle. Karniel and Mussa-Ivaldi showed that participants failed to learn alternating opposing force fields in sequential reaching tasks, suggesting that movement order alone is not sufficient to switch between opposing dynamics (26). A key difference from our task is that, in their task, each reach was triggered by the presentation of a new target only after the previous target had been acquired, likely encouraging participants to plan each movement separately. By contrast, our double-reach task encouraged participants to plan and execute multiple movement elements as a single action unit. Thus, sequence-level progression should not be viewed as an inevitable property of any movement sequence, but rather as a representational coordinate that may emerge when task structure encourages multiple movement elements to be organized as a single planned action.

### Action progress as a continuous memory dimension

A key limitation of Experiment 2 is that progression-dependent retrieval could, in principle, reflect a categorical distinction between the first and second reaches rather than a continuous coordinate. Experiment 3 addressed this issue using a button–reach–button paradigm in which the reach occurred at parametrically varied progression values within the sequence (Figs. 4a-b). This task was also designed to encourage sequence-level planning. As in Experiment 2, the temporal structure of the entire sequence was specified before movement onset: participants were informed in advance of the timing of all three movement elements. In addition, based on previous evidence that movements separated by intervals shorter than approximately 600 ms are likely to be planned in an integrated manner (23), we kept the intervals between movement elements below this range. Moreover, previous work has shown that actions involving different effectors can nevertheless be linked within a common sequence context (27, 28), supporting the idea that the button presses could provide a sequence context for the reach.

Participants learned opposing force fields more successfully when they were assigned to different progression values within the button–reach–button action than when both fields were assigned to the same progression value (Figs. 4f-g). However, in the button–reach–button task, relative progression was not the only potential cue: elapsed time and the visual position of the tick cue also differed between fields. If these cues alone were sufficient to separate opposing memories, then a simpler button–reach task should have supported a comparable degree of dual learning. Instead, learning was weaker in the button–reach task than in the button–reach–button task (Figs. 6e-g). This suggests that the critical factor was not merely the presence of temporal or visual cues, as also suggested by previous studies (29, 30), but whether the reach was defined by its relative position within the overall action sequence. This finding extends previous evidence that motor memories can be shaped by the relative timing between two movements (27), by showing that they can also depend on a movement’s relative position within an overall action sequence.

A further question is whether progression-dependent organization is specifically action-based or can also be defined by task-relevant events. One possibility is that the final button press in Group 1 acted not as an action element, but as an event that defined the sequence structure. If so, replacing this button press with a purely sensory cue should, in principle, still produce better memory separation than in Group 3. However, this interpretation is difficult to reconcile with previous evidence that purely visual contextual cues are relatively ineffective in separating opposing motor memories (29, 30). Thus, the difference between the button–reach–button and button–reach tasks is more consistent with the notion that the final button press, as an action element, helped define the reach’s relative position within the unfolding action sequence.

Nevertheless, progression through behavior may not be defined exclusively by overt action elements. Neural population activity has been shown to represent motor task-related information in a sequential and event-related manner (31), raising the possibility that behaviorally meaningful events, such as rewards, go cues, or goal-related signals, could also contribute to defining progression through a sequence. Further studies are therefore needed to determine whether progression-dependent memory expression is specific to action sequences or reflects a broader task-event coordinate.

The parametric design further allowed us to examine memory expression at untrained progression values, revealing graded compensation along the progression axis (Figs. 5a-b). Critically, individual differences in memory expression were predicted by how distinctly the two force fields were separated along progression, even when the nominal task conditions were identical (Fig. 5f). Together, these findings provide converging evidence that action progress serves not merely as a categorical contextual cue, but as an abstract, continuous coordinate for organizing motor memory within the structure of a planned action.

### Conceptual implications and future directions

Our findings extend the concept of motor abstraction from abstract representations of movement patterns to the abstract coordinates by which motor memories are organized and retrieved. Previous work has shown that abstract motor representations exist and can support the learning and generalization of novel movements beyond the conditions in which they were acquired (7,8). Yet motor memory has traditionally been described as a state-dependent process (13–18), in which memories are linked to physical variables such as position and velocity. This raises a fundamental question: how can memories learned through specific movements be flexibly reused when future actions differ in scale, timing, or sequence context? The present results identify action progress as an abstract organizing coordinate that complements physical-state representations. Importantly, this coordinate may provide a basis for organizing individual movement experiences directly into an abstract form, rather than reconstructing abstract motor memory only retrospectively from memories tied to separate physical states. This interpretation resonates with recent theoretical accounts of categorization, which propose that categorization is not an end stage of perception but a process that shapes signal processing from the outset through predictive, action-oriented abstractions (32). In a similar way, action progress may structure motor learning as an action unfolds, supporting both the construction and retrieval of abstract motor memory according to its functional position within an intended action. Thus, this study provides a framework for understanding how abstract motor memories support flexible motor control. Because the present study focused on motor adaptation, an important future direction is to determine whether similar progress-based coordinates also contribute to other forms of motor learning, such as de novo skill acquisition and reward-based learning.

The continuity revealed in Experiment 3 also has an important implication for flexible generalization. Unlike a categorical cue, a continuous progress coordinate allows learned outputs to vary smoothly with the relative position of a movement within a planned action. This continuity fits naturally with neural population dynamics accounts in which movement-related neural activity evolves along continuous, phase-like trajectories in neural state space (10, 11). However, evidence from rapid movement sequences suggests that motor cortex does not simply fuse successive elements into a single holistic dynamical cycle; instead, individual elements are generated through partly separate dynamics, with preparation for upcoming elements overlapping with ongoing execution (33, 34). Thus, sequence-level action progress may depend on higher-order population-level mechanisms that represent the structure and progression of an action sequence, consistent with evidence for ordinal-rank subspaces in prefrontal cortex and progression-dependent encoding geometries in posterior cortical populations (31, 35, 36). These representations may interact with cerebellar and basal ganglia circuits to organize multiple movement elements into appropriately timed and sequentially ordered actions, as dysfunction of these systems impairs the rhythmic and sequential organization of movement in cerebellar ataxia and Parkinson’s disease (37–41).

Finally, the present study bridges motor adaptation and sequence learning within a common framework. Motor adaptation and sequence learning have traditionally been treated as distinct paradigms in motor learning research (12). Sequence learning, in particular, has often been discussed in terms of hierarchical organization and chunking of movement elements (42–44). The present findings connect these traditions by showing that adaptation memories for a reach can depend on where that reach occurs within the structure of a planned action sequence. This perspective is important because everyday motor behavior rarely consists of isolated movements; rather, it is composed of action elements embedded within structured goals, sequences, and contexts. Future work should examine how progress-based coordinates are formed as action sequences are learned, and how hierarchical circuits involving motor cortical areas, higher-order cortical regions, and subcortical structures define planned action units and organize memory retrieval within them.

## Materials and Methods

### Experimental Design

This study tested whether motor memory is organized by action progress rather than by the specific movement states experienced during learning. We used three experiments. Experiment 1 examined whether motor learning generalized across movement scales in single reaching movements according to action progress. Experiment 2 tested whether distinct motor memories were associated with different progression ranges in a double-reach sequence. Experiment 3 examined whether motor memory was organized along a continuous sequence-level progression in a button–reach–button task. Participants were assigned to predefined groups, force-field directions were counterbalanced across participants, and motor memory was quantified using force output measured in error-clamp probe trials and learning traces in perturbation trials. Prespecified exclusion criteria were based on movement initiation, task timing, movement execution, and completion of the session.

### Participants

A total of 109 right-handed participants aged 18–35 years naive to the purpose of the study participated in the experiments as follows: Experiment 1 (Group 1: 8 (4 women); Group 2: 8 (4 women)), Experiment 2 (Group 1: 16 (8 women); Group 2: 17 (9 women); Group 3: 8 (4 women)), and Experiment 3 (Group 1: 18 (9 women); Group 2: 18 (10 women); Group 3: 16 (8 women)). Participants were compensated ¥2,000 per hour for their time.

All participants provided written informed consent after receiving a full explanation of the experimental procedures. All experimental procedures were conducted at the Center for Information and Neural Networks, National Institute of Information and Communications Technology (NICT), Japan. All study protocols followed the ethical standards of the Declaration of Helsinki and were approved by the Ethics Committee of NICT (Approval No. [N230082403]).

### Data and participant exclusion criteria

For each experiment, trials were excluded from analysis if any of the following criteria were met: (i) movement onset occurred before the go cue or too late (Experiment 1: more than 1 s after the trial start; Experiment 2: more than 1.2 s after the trial start); (ii) double movement occurred in which participants initiated an unintended pre-movement before the go cue and then executed the instructed movement again after the cue; (iii) the movement speed was too slow (< 0.05 m/s) in Experiment 3; or (iv) the first button press occurred outside the instructed temporal window in Experiment 3.

One participant discontinued Experiment 2 before completion and was therefore excluded from the analysis. To ensure that all participants performed the task in Experiment 3 with equivalent timing, a practice session was introduced before the main experiment (as described in the procedure of Experiment 3 below). Participants who could not achieve the predefined performance criterion during this practice session did not proceed to the main experiment.

Participants who took considerably longer and were unable to complete the task within the scheduled session time were excluded from analysis in Experiment 3 (two participants in Group 1 and two in Group 2).

### General setting

Participants performed reaching movements using their right hand while grasping the handle of a robotic manipulandum (PHANToM Premium 1.5 HF; SensAble Technologies) with a position sampled at 500 Hz. Visual stimuli, including a white cursor (3 mm in diameter) representing the handle position, were projected on a horizontal display positioned above the handle. This display prevented participants from directly viewing their hand and the manipulandum.

### Perturbations

Two types of perturbations were used in this study to elucidate how the motor system represents motor memory. In Experiment 1, a position-dependent force field was applied such that the direction of the force reversed between the first and second halves of the movement (20). The force acting on the handle was defined as follows:

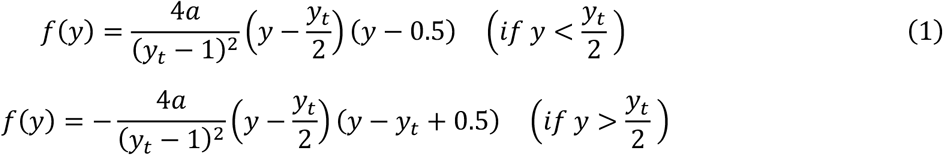

where y (cm) denotes the position along the movement direction, a is the maximum force gain, and *y_t_* (cm) represents the target distance. To prevent unwanted force generation before movement onset and after movement termination, a 0.5-cm buffer region was implemented at both the start and end of the movement range.

Experiments 2 and 3 utilized a velocity-dependent curl force field applied according to the following equation:

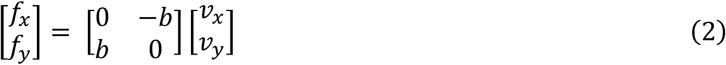

where *f_x_* and *f_y_* represent the forces applied to the handle (N), and *v_x_* and *v_y_* are the velocities of the handle (m/s) in the x and y directions, respectively. The viscosity coefficient b (N s/m) was positive for the CW force field and negative for the CCW field.

Error-clamp probe trials were occasionally introduced to quantify learning of these perturbations. During probe trials, the handle was constrained along a straight path between the start position and the target by a simulated damper and spring. The force applied to the clamp channel was measured, allowing measurement of the force retrieval to resist perturbations in each situation without experiencing the perturbation itself and kinematic error.

### Experiment 1

Experiment 1 examined the generalization of motor memory across different movement distances during single reaching movements.

#### Procedure

A gray circular target (4.5 cm in diameter) was presented 8 cm or 16 cm away from the start position in the straight-ahead direction. A few seconds after participants placed the handle at the start position, the target color changed to magenta, which served as the “go” signal. Participants were instructed to move the handle smoothly in a straight direction toward the target. The cursor was continuously visible during the reaching movement. The movement time was defined as the duration from when the handle left the start position to when it entered the target area. To maintain a natural and consistent movement speed, the target movement time was set to 350 ms for the 8 cm target and 500 ms for the 16 cm target. Participants were required to complete the movement within 0.75–1.25 times the target movement time. If the movement duration was shorter than this range, a “fast” warning message was displayed at the end of the trial; if the movement duration was longer, then a “slow” warning message was displayed.

Experiment 1 consisted of four sessions: a familiarization session to allow participants to adapt to the experimental environment, a baseline session to measure force output, a learning session, and a test session (Fig. 1a–b). In the familiarization session, the participants performed reaching movements to the two target distances (8 and 16 cm), which were randomly interleaved, with 20 trials per distance. In the baseline session, participants first performed four trials without force perturbation (two per target distance). This step was followed by 40 trials with the same structure as the familiarization session, with error-clamp trials randomly interleaved in 20 trials.

Participants were divided into two groups that differed in the distance at which they experienced the position-dependent S-shaped force perturbation during the learning and test sessions. Each group consisted of eight participants, guided by prior work (12, 19) and experimental design considerations, including counterbalancing of sex and movement direction. Group 1 learned the perturbation only at the 8 cm target (force gain a = ± 2 N; Fig. 1a), whereas Group 2 learned it only at the 16 cm target (force gain a = ± 3.5 N; Fig. 1b). The direction of the force field was counterbalanced across participants (four participants per direction). After ten null trials without force perturbation, the force magnitude gradually increased every five trials by ± 0.4 N for the 8 cm target in Group 1 and ± 0.7 N for the 16 cm target in Group 2. The full-strength perturbation was applied in the following 240 trials. Every tenth trial was a probe trial conducted at the same target distance as the perturbation trials to evaluate the progression of learning. No force perturbation was applied in the probe trials; instead, the handle was constrained to move along a straight path by a force channel (error clamp), with the lateral force exerted against the channel measured as retrieved motor memory. The 270 perturbation trials were divided into two sets (150 and 120 trials) with a 2-min rest period between sets.

After the learning session, a test session was conducted to evaluate retention and generalization. The test session consisted of three sets (130 trials each) with a 2 min rest period between sets. In each set, participants first performed 30 perturbation trials identical to those in the learning session, followed by 20 alternating cycles of four perturbation trials at the same distance as the learning session and one probe trial. For the probe trial, error-clamp trials were conducted at two target distances to examine the generalization of the learned response, which was quantified as the lateral force exerted against the channel clamp. The two target distances in the probe trial were presented in a pseudorandom order, with ten blocks for each target distance within a set. Participants thus performed a total of 754 trials.

### Experiment 2

Experiment 2 investigated whether the motor system encodes motor memory according to the progression of double reaches.

#### Procedure

Participants performed two types of double-reach movements: lead-in and follow-through (Fig. 2a). In both trial types, three targets were displayed at the beginning of each trial: the start location (4.5 cm in diameter; yellow), the central target (4.5 cm in diameter; gray), and the final target (4.5 cm in diameter; gray). In the lead-in trial, the handle was moved automatically to the start location. After maintaining the handle within the start position for approximately 1 s, the final target turned magenta to signal participants to initiate the movement. The participants then performed a lead-in movement from the start location to the central target (7.5 cm). This movement was always performed in the null force field. The central target served as the starting position for the subsequent main movement, which continued from the central target to the final target (7.5 cm). In the follow-through trial, participants performed the main movement from the start (yellow) to the central target (gray). This step was immediately followed by a follow-through unperturbed movement from the central target to the final target (7.5 cm). In both the lead-in and follow-through trials, the main movement was always directed toward the forward direction (0°). Movement directions were defined relative to the forward direction of the main movement (0°), with the rightward and leftward directions defined as positive and negative, respectively, and the backward direction as 180°.

Participants were instructed to perform the two sequential movements with as consistent a rhythm as possible, planning both movements as a single action, and stop briefly at the central target between the two movements. To control movement speed and avoid long stays at the central target, the duration from the go cue to the end of the movement was monitored. If this duration exceeded 1080 ms, participants received feedback indicating that their movement was too slow (“slow”) after the trial.

Experiment 2 consisted of three sessions for Group 1 and four sessions for Groups 2 and 3. These included the familiarization session, baseline session, learning session, and test session (not performed by Group 1) (Fig. 2d). Participants were assigned to one of three groups based on the type of trials they performed. Each group consisted of 16 participants, consistent with previous studies investigating lead-in and follow-through effects (e.g., 22, 23).

#### Group 1 (High interference group)

Group 1 was further divided into two subgroups: a lead-in group (Fig. 2b, left; n = 8) and a follow-through group (Fig. 2b, right; n = 8). Lead-in group participants performed four types of movement sequences: three lead-in trials, in which the main movement was preceded by a lead-in movement originating from the 180° (backward), +135° (backward-right), or −135° (backward-left) direction, and one follow-through trial, in which the main movement was followed by a follow-through movement directed toward 0° (forward). Follow-through group participants performed three follow-through trial types, in which the main movement was followed by a follow-through movement directed 0° (forward), +45° (rightward), or −45° (leftward), and one lead-in trial type, in which the main movement was preceded by a lead-in movement originating from 180° (backward).

During the familiarization session, participants experienced 10 of each trial type to become accustomed to the task. In the baseline session, participants performed 21 blocks, each consisting of four trial patterns presented in a randomized order. After the first block, every other block of the remaining 20 blocks included two probe trials where the corresponding normal trials, including the two forward movements, were replaced by trials performed under a channel clamp. The lead-in trial originating from the backward direction (180°) is referred to as Probe 1, and the follow-through trial directed forward (0°) is referred to as Probe 2.

In the learning session, participants performed movement sequences presented in a pseudorandom order. The learning sequences for the lead-in group included lead-in movements originating from +135° or −135°, while those for the follow-through group included follow-through movements directed toward +45° or −45°. Participants completed a total of 410 learning trials, with the first 10 trials performed without force perturbation for familiarization. The force-field coefficient (b) was set to +10 or −10, depending on the direction of the lead-in or follow-through movement (22, 23). Mapping between the movement and force-field directions was counterbalanced across participants. The 410 perturbation trials were divided into four sets (110, 100, 100, and 100 trials), with a 2 min rest period between sets.

The two opposing force fields used in this experiment are similar to those used in previous studies (22, 23). Although the fields were slightly weaker in magnitude, this design was selected to avoid disrupting the rhythmic pattern of the movements. We hypothesized that this environment would force participants to learn two conflicting motor memories within the same movement progression, thereby leading to interference.

#### Group 2 (Reduced interference group)

Group 2 was used to test whether distinct motor memories are formed in different progression ranges of a double-reach movement. Each participant was assigned to either the left or the right side. Participants performed four types of movement sequences (Fig. 2c): two follow-through trials, in which the main movement was followed by a follow-through movement directed toward 0° (forward) or toward the participant-assigned side (−45° for left, +45° for right), and two lead-in trials, in which the main movement was preceded by a lead-in movement originating from 0° (forward) or from the backward direction on the same assigned side (−135° for left, +135° for right).

The familiarization and baseline session procedures for Group 2 were the same as those for Group 1, except that the four trial types reflected the above-described movement patterns. In the learning session, lead-in trials originating from the left or right side (±135°) and follow-through trials directed toward the left or right side (±45°) were presented in a randomized order. The force-field coefficient (b) was set to +10 or −10, depending on the trial type (lead-in or follow-through). The correspondence among trial types, applied field direction, and participant-assigned side was counterbalanced across participants.

In the subsequent test session, participants completed two test sets with a 2-min rest period between sets. For each set, participants performed 20 perturbation trials under each force-field condition, followed by repeated test blocks designed to assess memory retrieval. Each test block consisted of six perturbation trials (three per movement sequence) and one probe trial (Probe 1 or Probe 2), with trial order randomized within each block. Each set included 20 such blocks. This design allowed us to quantify the retrieved motor memory for each reaching movement using the probe trials. Participants completed a total of 854 trials across all sessions.

#### Group 3 (Single motor memory group)

For this group, the lead-in trials originating from the left or right side (±135°) and the follow-through trials directed toward the left or right side (±45°) used for Group 2 were both replaced with single-reach movements performed at the corresponding main-movement locations. A velocity-dependent curl field with a coefficient (b) of +10 or −10 was applied for the learning and test sessions (Fig. 3h inset), with only one field direction assigned to each participant (counterbalanced across participants). All other procedures were identical to those for Group 2. This configuration allowed us to examine how a single motor memory acquired in a single-reach movement is expressed when participants perform the double-reach task.

### Experiment 3

Experiment 3 established whether motor memory is organized along a continuous sequence-level progression.

#### Procedure

Participants performed a sequence comprising three consecutive actions: pressing a keyboard key with the left index finger (B_1_), smoothly reaching forward with the right hand toward a target located 4 cm away (R), and pressing another key with the left index finger (B_2_). Participants were instructed to pre-plan the timing of these actions and execute them as a continuous, well-timed sequence.

A horizontal progress bar displayed on a screen indicated the temporal structure of the task (Fig. 5b). Five vertical reference lines were drawn on the bar, corresponding to −750 ms, −375 ms, 0 ms (the timing of the first button press), the timing of the reach, and the timing of the second button press. The start position for the reach was located just below the line corresponding to the reach time. After participants held the cursor at the start position for approximately 1 s, the progress bar appeared for 1 s to allow them to understand the temporal structure of the upcoming trial. The bar then began to fill from left to right in pink, starting from the leftmost line. The interval of 375 ms corresponded to a horizontal distance of 3.9 cm on the display. When the bar had filled with pink to the third line, it stopped, and the participants were instructed to imagine its continuation and perform the movement such that the pink bar would overlap with each reference line at the appropriate timing. During the practice session, small feedback markers were presented above the progress bar in real time at the timing of each button press and the onset of the reach (defined as 8 mm of movement). Participants were encouraged to adjust their performance so that these feedback markers aligned with the reference lines during the trials. If the peak velocity of the reaching movement deviated from the prescribed range of 0.3–0.4 m/s, the word “slow” or “fast” was displayed as feedback on the screen after the trial. A trial was considered successful if the timing error for each action was within 150 ms of the target timing.

Participants were divided into three groups according to the type of movement they performed. Each group consisted of 16 participants, with the sample size matched to that used in Experiment 2 to maintain consistency across experiments. All participants completed four experimental sessions: a practice session, a baseline session, a learning session, and a test session. The practice session consisted of three training sets that were designed to familiarize participants with the temporal structure of the movement sequence. Training set trials were considered successful if the timing error of each action in the sequence was within ±180 ms of the instructed timing. In the first set, participants practiced aligning only the button-press actions (B_1_ and B_2_) with the corresponding timing cues of the progress bar. In the second set, participants practiced aligning the reaching movement with the appropriate timing. In the last set, participants synchronized the button-press and reaching movements with the specified temporal structure. The final set was completed when participants successfully performed either eight consecutive correct trials or six consecutive correct trials after a total of 100 attempts, whichever occurred first. The temporal structure of the task was randomly selected from the set of structures used in the main experiment. Across training sessions, the participants learned to internally predict the progression of the invisible bar and complete the B_1_–R–B_2_ sequence so that each action aligned with equally spaced segments on this timeline, thus pre-planning the entire sequence before initiating the first movement.

The second button press (B_2_) was omitted for Group 3 participants, who only performed the first button press followed by the reaching movement (Fig. 6a) while maintaining the temporal constraints described above. Following the practice session, timing feedback was only provided after completing the sequence to minimize online timing corrections.

#### Group 1 (Reduced interference group)

For Group 1, the interval between the two button presses (B_1_–B_2_) was fixed at 750 ms throughout the experiment (Fig. 4d). Participants performed three types of trials with this fixed interval that differed in the timing of the reaching movement (R): 250 ms (early), 375 ms (middle), or 500 ms (late) after the first button press. For the baseline session, participants performed a total of 44 trials with the 375-ms middle condition. The first four trials were performed in the null field. These trials were followed by alternating probe trials with error clamp trials and normal trials to measure baseline force output during reaching.

For the learning session (Fig. 4c), participants completed a total of 440 trials. Early and late conditions were presented in a pseudorandom order, such that the two conditions alternated across trials without forming a fixed sequence. The first ten trials were conducted in the null field to allow participants to adapt to the task before introducing the perturbation. Subsequently, a velocity-dependent curl force field was applied with the force field coefficient (B) set to +8 or −8, counterbalanced across participants. This force-field magnitude was weaker than that used in previous studies examining conflicting dynamic perturbations (22, 23, 28, 29), in order to minimize disruption of movement timing and preserve the temporal structure of the task. Participants took short breaks of 1–2 min every 110 trials to prevent fatigue.

For the test session, participants completed three sets with a 2 min rest period between sets. Each set began with 10 perturbation trials performed under the same conditions as in the learning session, followed by repeated blocks consisting of six perturbation trials (three per field direction) and one probe trial (Early, Middle, or Late). An error-clamp channel was imposed in the probe trials to measure the lateral force, as in Experiments 1 and 2. This block was repeated 12 times (four repetitions per timing condition) for the first and second sets and six times for the third set (two repetitions per timing condition).

#### Group 2 (High interference group)

Group 2 performed three types of trials that differed in the interval between the button presses: 500 ms (Early), 750 ms (Middle), and 1000 ms (Late). The progression value of the reaching movement was always centered between the two button presses (B_1_–B_2_). This design allowed variation in the absolute timing of the sequence while fixing reach at the same progression value, thus isolating the effect of progression on motor memory formation (Fig. 4e). Because the two force fields were assigned to the same progression value, this configuration enabled examination of the extent to which learning interference arises when progression-based separation is unavailable. This procedure was identical to that for Group 1, except that the three sequences (Early, Middle, and Late) were defined by the B_1_–B_2_ interval rather than by the progression between two button presses.

#### Group 3 (absolute-time group)

The final button press (B_2_) was omitted for Group 3, and the interval between the first button press (B_1_) and the reach (R) was set to one of three values: 250 ms (early), 375 ms (Middle), or 500 ms (Late). The direction of the velocity-dependent curl field was determined according to the absolute B_1_–R interval (Fig. 6a). This design eliminated progression-based differences between the conditions and allowed examination of how absolute timing from a preceding movement contributes to the formation of motor memory. This procedure was identical to that for Groups 1 and 2, except that the three sequences (Early, Middle, and Late) were defined by the absolute B_1_–R interval rather than by progression within the movement sequence.

### Data analysis

The handle position and force data were sampled at a rate of 500 Hz and filtered using the 4th-order zero-lag Butterworth filter with a cut-off frequency of 10 Hz. Data analysis was performed using MATLAB v.2020b (Mathworks). For all analyses, the lateral force profile was defined as the lateral forces exerted in the probe trials, subtracting the average baseline force from the clamp channel in the baseline session.

#### Data analysis for Experiment 1

Movement onset was determined as the time when the handle velocity reached 5% of the peak velocity. The relationship between the lateral forces and applied forces was analyzed by aligning all trials at the movement onset and averaging them across repetitions. Because the force field direction was counterbalanced across participants, force data were flipped as necessary so that the perturbation direction in the first half of the movement corresponded to the negative direction across participants. The direction of the retrieved lateral force was quantified in the early and later parts of the movement. The maximum value of the lateral force exerted in the direction opposing the perturbation within each range of the progression was taken as an index of motor memory expressed in the appropriate compensatory direction. To assess force output at comparable spatial locations across groups, the integrated lateral force was computed over a 6–7.5-cm region (Fig. 1e inset). These measures enabled the determination of whether learned motor memory was expressed in a state- or progression-dependent manner.

To examine whether the force profiles could be explained by memory tied to absolute elapsed time, probe trials were divided within each participant into fast and slow subsets (Supplemental Figure 1). Specifically, the eight trials with the shortest and longest times to reach the midpoint distance were labeled fast and slow trials, respectively. Force profiles were then averaged within each subset. To quantify temporal scaling, the fast-trial force profile was temporally scaled and correlated with the slow-trial force profile, and the scaling factor that maximized this correlation was identified for each participant.

To examine whether the force profiles could be explained by acceleration-dependent representations, force and acceleration profiles were plotted as a function of position (Supplemental Figure 2). For each group and reaching distance, the positions of the maximum and minimum peaks were identified for both force and acceleration profiles and compared using paired t-tests.

#### Data analysis for Experiment 2

The first movement onset was defined as the time when the handle velocity reached 5% of its peak value, while the second movement onset was defined as the time when the handle velocity reached 10% of its peak value. Different velocity thresholds were used because the first movement started from rest, whereas the second movement was initiated immediately after completion of the first movement. This step made it difficult to accurately detect its onset with a lower threshold and could falsely identify small residual movements as movement onset. To quantify learning progress during the learning session, the absolute lateral deviation of the main movement from the straight path was measured within the progression range in which the perturbation was applied. For each block, this deviation was compared with the mean deviation observed in the initial null trials at the beginning of the learning session, with the difference defined as the error.

Two complementary metrics were used to quantify error reduction across groups. First, overall facilitation was assessed using an error reduction ratio, defined as 1 minus the ratio of the mean lateral deviation (E) in the last ten trials (i.e., the last block) to that in the first perturbed ten trials (i.e., the first block) of the learning session (1 − *E_last_*/*E_first_*)). This metric quantified the extent to which error decreased over the learning period relative to initial performance. Second, the temporal profile of error reduction was characterized by fitting an exponential decay function to the trial-by-trial time course of lateral deviation as follows: *y* = *Ae*^−*Bx*^ + *C*, where y denotes lateral deviation, x denotes the block number, and B represents the error decay rate. This parameter of the fitted curve was extracted as an index of learning speed.

During the test session, participants performed two types of probe trials. The exerted lateral force was time-aligned to the movement onset for the first and second movements and then averaged across trials. Because the direction of the force field was counterbalanced across participants, force data were flipped as necessary to align the direction of perturbation across participants. To quantify the direction of the retrieved motor memory, the mean force exerted during the pre-movement period was subtracted from the force profile in the second reach, as this force primarily reflected the residual effect of the preceding movement. The peak value of the resulting force trace was taken as the magnitude of the retrieved motor memory (Fig. 3e inset).

We calculated the ratio between the peak force of the second reach and that of the first reach in the same movement to examine whether the retrieved motor memory reversed in a movement depending on the progression of the sequence. We also calculated the ratio between the peak force of the second reach in Probe 1 and the first reach in Probe 2 to examine whether the retrieved motor memory differed with progression of the sequence, despite being expressed in the same kinematic state. These ratios were compared between Groups 2 and 3 to assess motor memory retrieval during the double reach with a clamp channel.

#### Data analysis for Experiment 3

The first movement onset was defined as the time when the handle’s velocity reached 5% of its peak value. To quantify learning progress during the learning session, the deviation from the straight path was measured as in Experiment 2. The difference between this deviation and that observed in the baseline trials was defined as the deviation value. The error reduction ratio and decay parameter were quantified as in Experiment 2.

During the test session, participants performed three types of probe trials. The learning responses were time-aligned to the movement onset. Because the direction of the force field was counterbalanced across participants, force data were flipped as necessary to align the direction of perturbation across participants. Retrieved motor memory was quantified as the compensatory force impulse, defined as the impulse of the lateral force from movement onset to peak velocity. This value was normalized by the impulse of the ideal compensatory force, calculated as the hand velocity multiplied by the force-field constant b and expressed as a percentage (Fig. 5b). The compensatory force impulse size was computed separately for each probe condition (Early, Middle, and Late) for each participant. For summary analysis of overall motor memory retrieval, the size of the compensatory force from the two trained conditions (Early and Late) was averaged after aligning their signs so that positive values indicated forces in the appropriate compensatory direction. This averaged value was defined as the compensation ratio.

#### Model in Experiment 3

To test whether learning could be explained by how distinctly the two opposing force-field contexts were represented, we modeled the degree of separation of motor memories along three potential representational dimensions (Figs. 5d-e): progression, the absolute time between button press and reach onset (B_1_–R), and the absolute time between reach offset and the subsequent button press (R–B_2_).

Each dimension was discretized into small bins. For each bin, the occurrence of the two force-field types (CW and CCW) was accumulated. For each trial, a value of +1 was assigned for CW and −1 for CCW. The separation index for each dimension was then calculated as the sum of the absolute accumulated values across all bins as follows:

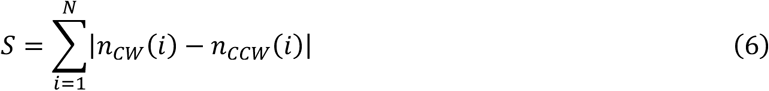

where N denotes the number of bins (100 bins for progression, with Δ progression = 0.01 × 2π; 100 bins for the absolute time dimensions, with Δt = 10 ms). A larger separation index indicates greater contextual differentiation between the two force fields within that representational dimension. Next, we tested whether these separation indices could predict the compensation ratio using linear regression analyses (without an intercept) between the separation index and the compensation ratio for each participant. The model was first applied independently for the three dimensions (progression, B_1_–R, and R–B_2_) and then extended to combined models (e.g., progression + B_1_–R) to evaluate which combination best explained across-subject differences in the observed compensation ratio, as assessed by the AIC.

### Statistical analysis

Unless otherwise specified, two-sided paired t-tests were used for within-subject comparisons and two-sided unpaired t-tests were used for between-group comparisons. For analyses of ratio-based metrics and the error decay rate, which may not follow a normal distribution, nonparametric tests were used. We performed one-sample comparisons against zero using the Wilcoxon signed-rank test and between-group comparisons using the Wilcoxon rank-sum test (Mann–Whitney U test). For Experiment 3, comparisons across the three groups used different statistical approaches depending on the metric. Group differences in the averaged compensation ratio, which served as a summary measure of overall expression performance, were assessed using one-way ANOVA. The Kruskal–Wallis test was used for other metrics in Experiment 3 that were not assumed to follow a normal distribution. In cases where multiple comparisons were performed, the p-values were adjusted using the Holm correction.

## Acknowledgments

We thank members of the Hirashima Laboratory for their helpful comments and suggestions, as well as for coordinating the experiments. This study was supported by the following grants: Japan Society for the Promotion of Science (JSPS) Research Fellowships for Young Scientists (24KJ2221 to YM).

## Author contributions

Y.M. and M.H. conceptualized the study. Y.M., K.S. and M.H. developed the methodology. Y.M., K.S. and M.H. performed the formal analysis. Y.M. conducted the investigation. Y.M. wrote the original draft. Y.M., K.S. and M.H. reviewed and edited the manuscript. M.H. supervised the project.

## Declaration of generative AI and AI-assisted technologies in the writing process

During the preparation of this manuscript, the authors used ChatGPT-5.5 model developed by OpenAI to improve readability, language, and clarity of the text. After using this tool, the authors reviewed and edited the content as needed. The authors take full responsibility for the content of the manuscript.

## Data availability

The data and code supporting the findings of this study are available from the authors upon reasonable request.

## Supplemental Figures

**Supplemental Figure 1.**
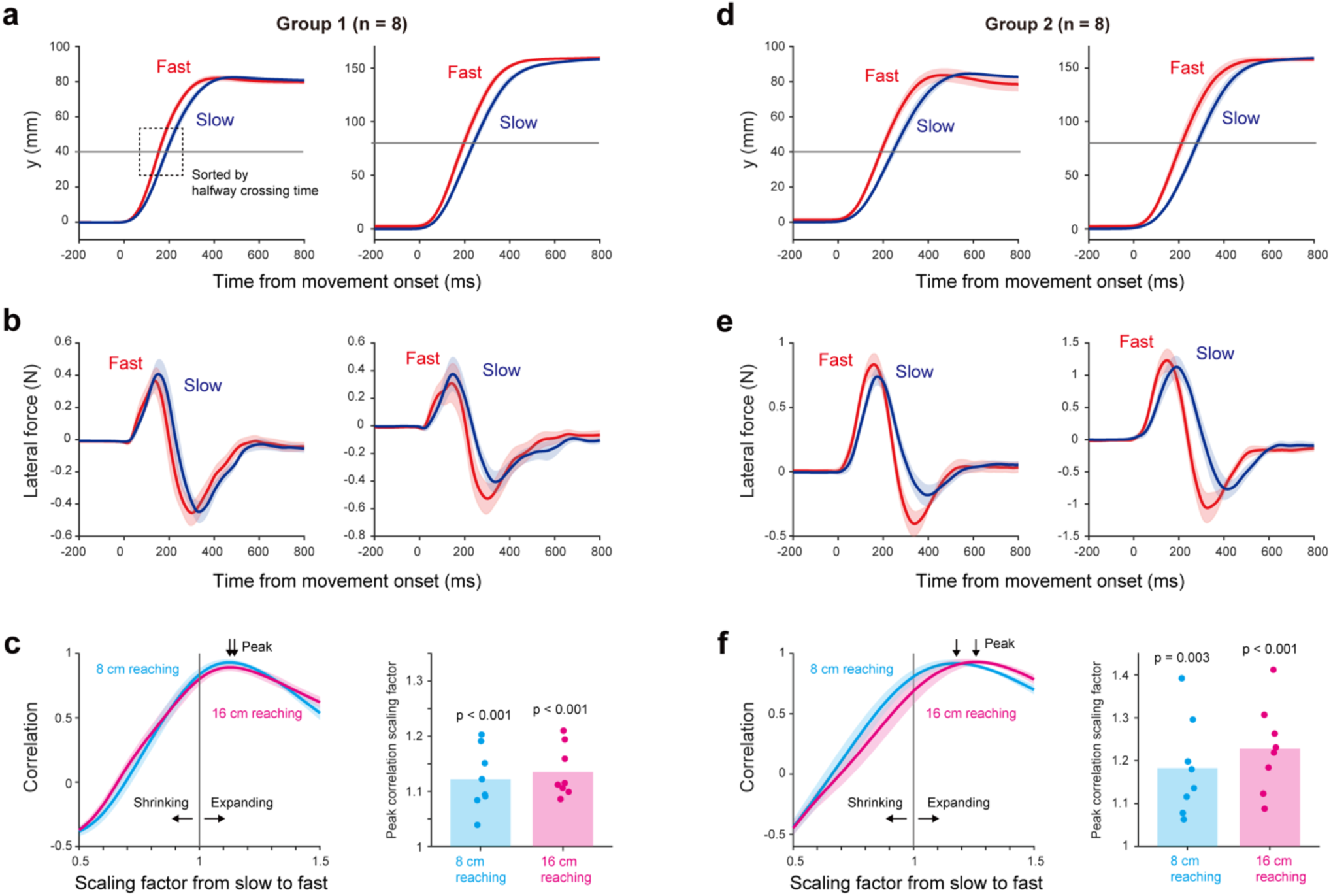
Absolute-time-dependent representation cannot simply explain the generalization in experiment 1. To test whether the probe-trial force profiles in Experiment 1 could be accounted for by memory tied to absolute elapsed time, trials were divided within each participant into fast and slow subsets. Specifically, the eight trials with the shortest and longest times to reach the midpoint distance—4 cm in the 8-cm condition and 8 cm in the 16-cm condition—were labeled fast and slow trials, respectively. The force profiles were then averaged within each subset. **a–b,** Position (a) and force (b) profiles for the fast and slow trial subsets in Group 1. If force generation were tied to absolute elapsed time, the temporal evolution of the force profiles should not differ between the fast and slow trial subsets. However, the force profiles instead appeared to scale with movement duration. **c,** Scaling analysis of force profiles. To quantify temporal scaling, the fast-trial force profile was temporally scaled and correlated with the slow-trial force profile. The left panel shows the correlation as a function of the scaling factor, and the right panel shows the scaling factor that maximized the correlation for each participant. The peak scaling factor was significantly greater than 1 (one-sample t-test; 8-cm reaching: t(7) = 6.116, p < 0.001; 16-cm reaching: t(7) = 8.216, p < 0.001), indicating that the force profiles were stretched in time rather than generated according to a fixed absolute-time pattern. d–f, Same analysis for Group 2. Position (d) and force (e) profiles for the fast and slow trial subsets, and the temporal scaling analysis (f). The peak-correlation scaling factor was significantly greater than 1 (one-sample t-test; 8-cm reaching: t(7) = 4.580, p = 0.003; 16-cm reaching: t(7) = 6.284, p < 0.001). These results suggest that force was not generated based on absolute elapsed time, supporting the idea that the memory was retrieved according to relative temporal progression within the movement.

**Supplemental Figure 2.**
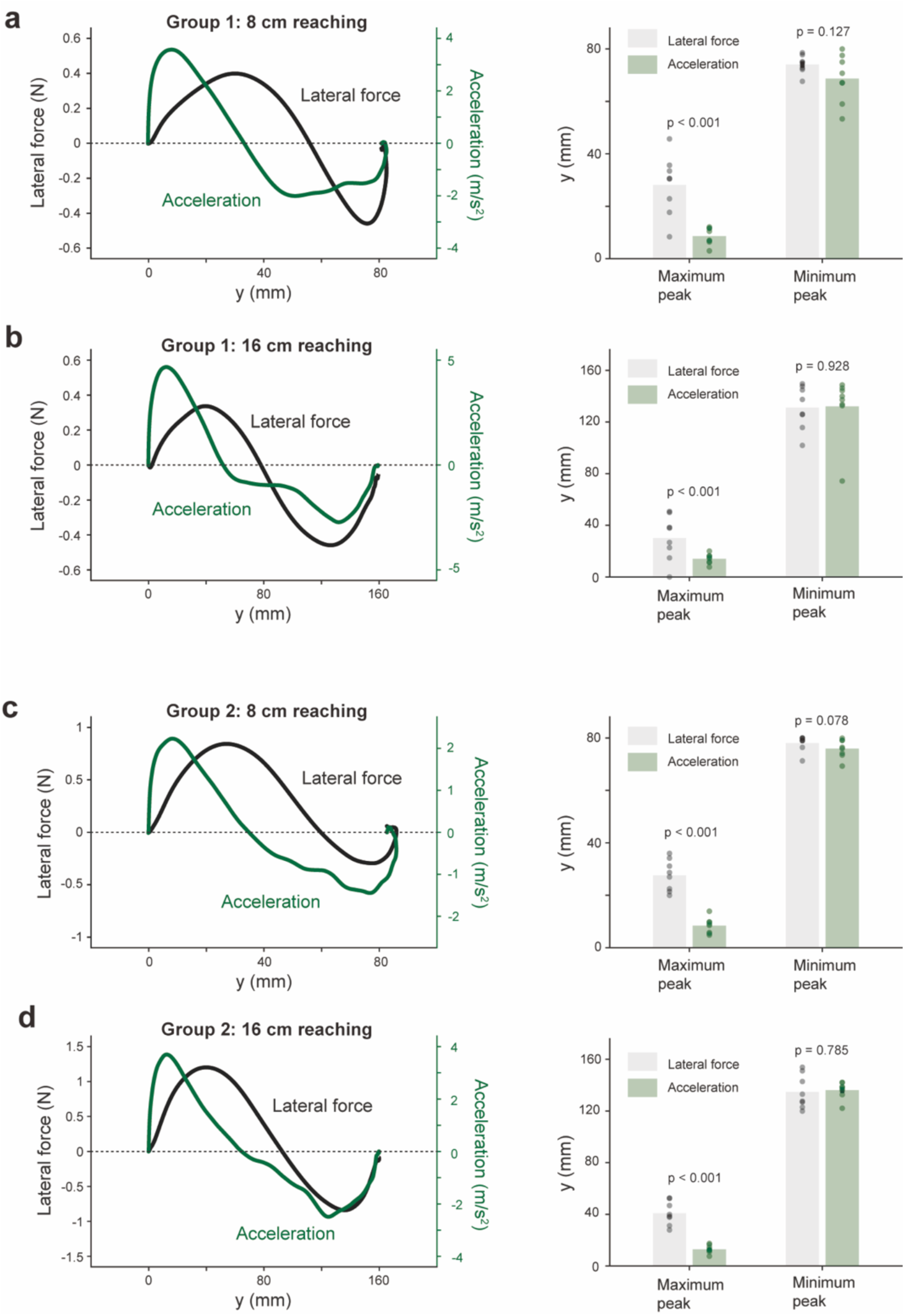
Acceleration-dependent representation cannot simply explain the generalization in experiment 1. **a–d,** Relationship between force, acceleration, and position. For each condition, the left panel shows force and acceleration profiles plotted as a function of position, and the right panel shows the positions of the maximum and minimum peaks. Panels show Group 1 during 8-cm reaching (**a**) and 16-cm reaching (**b**), and Group 2 during 8-cm reaching (**c**) and 16-cm reaching (**d**). The position of the maximum acceleration peak differed significantly from that of the maximum force peak in all conditions (paired t-test; Group 1, 8-cm: t(7) = −6.005, p < 0.001; Group 1, 16-cm: t(7) = −2.590, p = 0.036; Group 2, 8-cm: t(7) = −8.838, p < 0.001; Group 2, 16-cm: t(7) = −8.522, p < 0.001), whereas the minimum peak positions did not differ significantly (Group 1, 8-cm: t(7) = −1.731, p = 0.127; Group 1, 16-cm: t(7) = 0.093, p = 0.928; Group 2, 8-cm: t(7) = −2.060, p = 0.078; Group 2, 16-cm: t(7) = 0.284, p = 0.785). These results indicate that the force profiles are unlikely to be explained simply by acceleration-dependent representations.

**Supplemental Figure 3.**
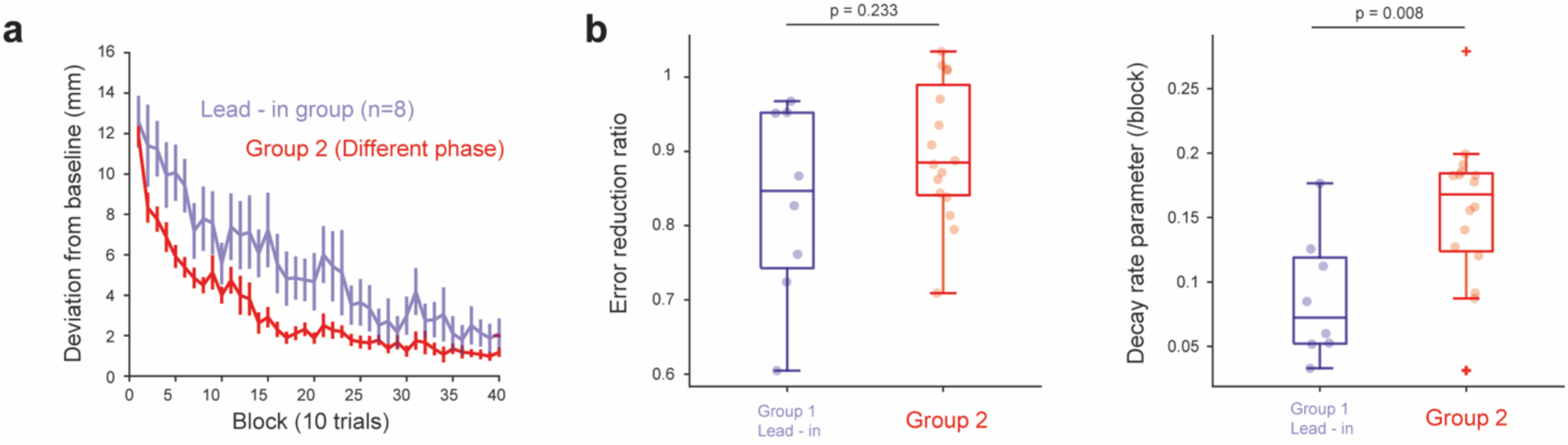
Comparison of learning traces between the lead-in subgroup (Group1) and Group 2 in Experiment 2. This analysis restricted the comparison to participants in the lead-in subgroup of Group 1, which is expected to exhibit facilitated learning than the follow-through subgroup based on previous studies (Howard et al., 2012, 2015). **a,** Learning traces for the two groups. **b,** The error reduction ratio did not differ significantly between groups (left; Mann–Whitney U test, z = 1.194, p = 0.232). However, the error decay rate differed significantly between groups (right; Mann–Whitney U test, z = 2.664, p = 0.008). Notably, Group 2 exhibited significantly facilitated learning compared with the lead-in subgroup of Group 1.

**Supplemental Figure 4.**
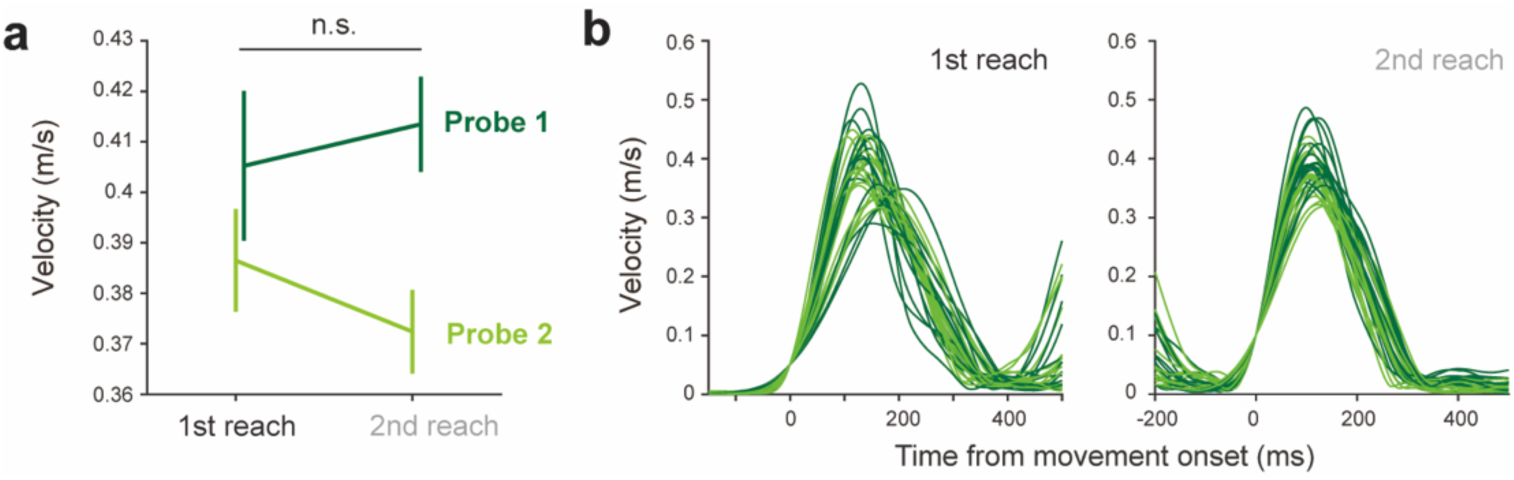
Similar temporal patterns of velocity between the first and second reaches in the Experiment 2 probe trials. **a,** Peak velocity in the Group 2 probe trials. Two-way repeated-measures ANOVA revealed no significant difference in velocity between the first and second reaches (F(1,15)=0.163, p=0.692). **b,** Temporal profile of velocity. After completing the first reach, the velocity decreased to nearly 0. The temporal profiles of the first and second reaches were highly similar.

**Supplemental Figure 5.**
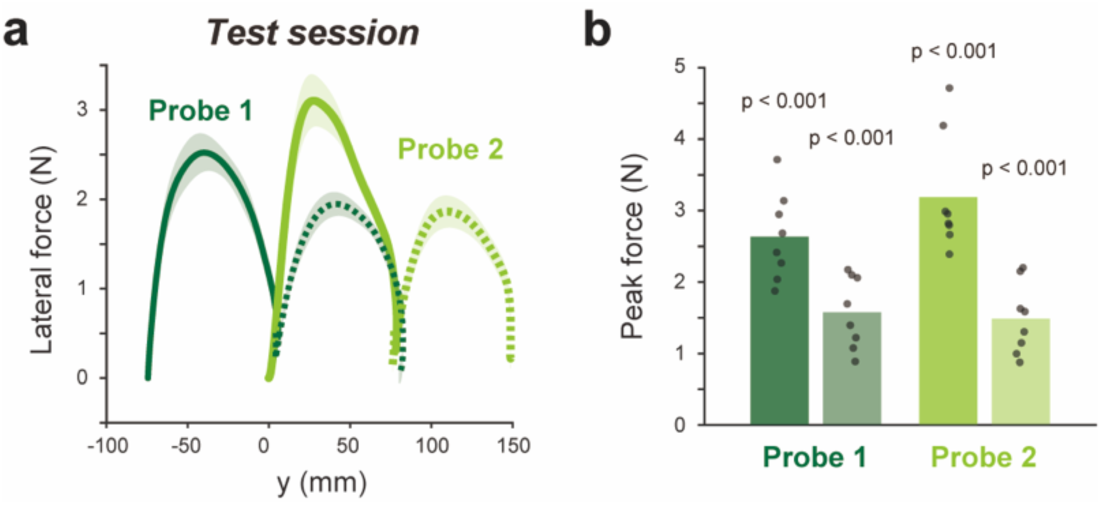
Learning response during double forward reaches for Group 3 in Experiment 2. Group 3 (Control) participants learned a force field in the single reach. We then examined how motor memory was retrieved during a double forward reach. **a,** Exerted forces in the double reaches. **b,** Peak forces observed in each reach. The peak forces were all significantly positive in Probe 1 (one-sample t-test; first reach: t(7)=12.153, p < 0.001; second reach: t(7)=8.899, p < 0.001) and in Probe 2 (one-sample t-test; first reach: t(7)=11.095, p < 0.001; second reach: t(7)= 8.439, p < 0.001).

## References

1. Tuller, B. & Kelso, J. A. S. The timing of articulatory gestures: Evidence for relational invariants. Journal of the Acoustical Society of America 76, 1030–1036 (1984).

2. Carter, M. C. & Shapiro, D. C. Control of sequential movements: Evidence for generalized motor programs. J. Neurophysiol. 52, 787–796 (1984).

3. van der Graaff, E., Hoozemans, M. M. J. M., Nijhoff, M., Davidson, M., Hoezen, M. & Veeger, D. H. E. J. Timing of peak pelvis and thorax rotation velocity in baseball pitching. J. Phys. Fitness Sports Med. 7, 269–277 (2018).

4. Viviani, P. & Terzuolo, C. Space–time invariance in learned motor skills. in Tutorials in Motor Behavior (eds Stelmach, G. E. & Requin, J.) 525–533 (North-Holland, 1980).

5. Shapiro, D. C., Zernicke, R. F. & Gregor, R. J. Evidence for generalized motor programs using gait pattern analysis. J. Mot. Behav. 13, 33–47 (1981).

6. Schmidt, R. A. A schema theory of discrete motor skill learning. Psychol. Rev. 82, 225–260 (1975).

7. Sun, Z., Xie, Z. & McDougle, S. D. Motor abstraction training generalizes to the refinement of specific movement patterns. bioRxiv 10.64898/2026.05.05.722946 (2026).

8. Tian, L. Y. et al. Neural representation of action symbols in primate frontal cortex. Nature 654, 152–162 (2026).

9. Ijspeert, A. J., Nakanishi, J., Hoffmann, H., Pastor, P. & Schaal, S. Dynamical movement primitives: Learning attractor models for motor behaviors. Neural Comput. 25, 328–373 (2013).

10. Churchland, M. M. et al. Neural population dynamics during reaching. Nature 487, 51–56 (2012).

11. Shenoy, K. V., Sahani, M. & Churchland, M. M. Cortical control of arm movements: a dynamical systems perspective. Annu. Rev. Neurosci. 36, 337–359 (2013).

12. Krakauer, J. W., Hadjiosif, A. M., Xu, J., Wong, A. L. & Haith, A. M. Motor learning. Compr. Physiol. 9, 613–663 (2019).

13. Sing, G. C., Joiner, W. M., Nanayakkara, T., Brayanov, J. B. & Smith, M. A. Primitives for motor adaptation reflect correlated neural tuning to position and velocity. Neuron 64, 575–589 (2009).

14. Hwang, E. J., Smith, M. A. & Shadmehr, R. Adaptation and generalization in acceleration-dependent force fields. Exp. Brain Res. 169, 496–506 (2006).

15. Conditt, M. A., Gandolfo, F. & Mussa-Ivaldi, F. A. The motor system does not learn the dynamics of the arm by rote memorization of past experience. J. Neurophysiol. 78, 554–560 (1997).

16. Conditt, M. A. & Mussa-Ivaldi, F. A. Central representation of time during motor learning. Proc. Natl Acad. Sci. USA 96, 11625–11630 (1999).

17. Diedrichsen, J., Criscimagna-Hemminger, S. E. & Shadmehr, R. Dissociating timing and coordination as functions of the cerebellum. J. Neurosci. 27, 6291–6301 (2007).

18. Shadmehr, R. & Mussa-Ivaldi, F. A. Adaptive representation of dynamics during learning of a motor task. J. Neurosci. 14, 3208–3224 (1994).

19. Thoroughman, K. A. & Shadmehr, R. Learning of action through adaptive combination of motor primitives. Nature 407, 742–747 (2000).

20. Albert, S. T. et al. Postural control of arm and fingers through integration of movement commands. eLife 9, e52507 (2020).

21. Scheidt, R. A. et al. Persistence of motor adaptation during constrained, multi-joint arm movements. J. Neurophysiol. 84, 853–862 (2000).

22. Howard, S., Ingram, J. N., Franklin, D. W. & Wolpert, D. M. Gone in 0.6 seconds: The encoding of motor memories depends on recent sensorimotor states. J. Neurosci. 32, 12756–12768 (2012).

23. Howard, S., Wolpert, D. M. & Franklin, D. W. The value of the follow-through derives from motor learning depending on future actions. Curr. Biol. 25, 397–401 (2015).

24. Haith, A. M. & Krakauer, J. W. Model-based and model-free mechanisms of human motor learning. in Advances in Experimental Medicine and Biology 782, 1–21 (2013).

25. Joiner, W. M., Ajayi, O., Sing, G. C. & Smith, M. A. Linear hypergeneralization of learned dynamics across movement speeds reveals anisotropic, gain-encoding primitives for motor adaptation. J. Neurophysiol. 105, 45–59 (2011).

26. Karniel, A. & Mussa-Ivaldi, F. A. Sequence, time, or state representation: how does the motor control system adapt to variable environments? Biol. Cybern. 89, 10–21 (2003).

27. Howard, I. S., Franklin, S. & Franklin, D. W. Kernels of motor memory formation: temporal generalization in bimanual adaptation. J. Neurosci. 44, e0359242024 (2024).

28. Gippert, M. et al. Prior movement of one arm facilitates motor adaptation in the other. J. Neurosci. 43, 4341–4351 (2023).

29. Howard, I. S., Wolpert, D. M. & Franklin, D. W. The effect of contextual cues on the encoding of motor memories. J. Neurophysiol. 109, 2632–2644 (2013).

30. Cothros, N., Wong, J. & Gribble, P. L. Visual cues signaling object grasp reduce interference in motor learning. J. Neurophysiol. 102, 2112–2120 (2009).

31. Koay, S. A. et al. Sequential and efficient neural-population coding of complex task information. Neuron 110, 328–349.e11 (2022).

32. Barrett, L. F. & Miller, E. K. Categorization is ‘baked’ into the brain. Nat. Rev. Neurosci. 27, 435–456 (2026).

33. Zimnik, A. J. & Churchland, M. M. Independent generation of sequence elements by motor cortex. Nat. Neurosci. 24, 412–424 (2021).

34. Schimel, M., Kao, T.-C. & Hennequin, G. When and why does motor preparation arise in recurrent neural network models of motor control? eLife 12, RP89131 (2024).

35. Xie, Y. et al. Geometry of sequence working memory in macaque prefrontal cortex. Science 375, 632–639 (2022).

36. Ding, N. Sequence chunking through neural encoding of ordinal positions. Trends Cogn. Sci. 29, 641–654 (2025).

37. Holmes, G. The symptoms of acute cerebellar injuries due to gunshot injuries. Brain 40, 461–535 (1917).

38. McLennan, E., Nakano, K., Tyler, H. R. & Schwab, R. S. Micrographia in Parkinson’s disease. J. Neurol. Sci. 15, 141–152 (1972).

39. Louis, E. D. et al. High width variability during spiral drawing: Further evidence of cerebellar dysfunction in essential tremor. Cerebellum 11, 872–879 (2012).

40. Letanneux, J., Danna, J., Velay, J.-L., Viallet, F. & Pinto, S. From micrographia to Parkinson’s disease dysgraphia. Mov. Disord. 29, 1467–1475 (2014).

41. Fujisawa, Y. & Okajima, Y. Characteristics of handwriting of people with cerebellar ataxia: Three-dimensional movement analysis of the pen tip, finger, and wrist. Phys. Ther. 95, 1547–1558 (2015).

42. Yokoi, A. & Diedrichsen, J. Neural organization of hierarchical motor sequence representations in the human neocortex. Neuron 103, 1178–1190.e7 (2019).

43. Diedrichsen, J. & Kornysheva, K. Motor skill learning between selection and execution. Trends Cogn. Sci. 19, 227–233 (2015).

44. Graybiel, A. M. The basal ganglia and chunking of action repertoires. Neurobiology of Learning and Memory 70, 119–136 (1998).

